# UniSpec: A Deep Learning Approach for Predicting Energy-Sensitive Peptide Tandem Mass Spectra and Generating Proteomics-Wide In-Silico Spectral Libraries

**DOI:** 10.1101/2023.06.14.544947

**Authors:** Joel Lapin, Xinjian Yan, Qian Dong

## Abstract

In this report, we present UniSpec, an attention-based deep neural network designed to predict complete collision-induced fragmentation of tryptic peptides, aimed at enhancing peptide and protein identification in shotgun proteomics studies. We preprocessed spectral data from peptide tandem mass spectral libraries, compiled by the National Institute of Standards and Technology (NIST), utilizing a data approach tailored for model development, resulting in high-quality, energy-consistent spectral datasets. By analyzing all the annotated fragment ions present in these libraries, we constructed an extensive peptide fragment dictionary containing 7919 isotopic ions from sequence ions, neutral loss, internal, iminium, and amino acid fragment ions. The streamlined dictionary-based spectral training data enables UniSpec to efficiently learn the complex intensity patterns of various product ions, resulting in reliable spectral predictions for a wide range of unmodified and modified peptides. We evaluated the model’s accuracy by comparing its performance across training and testing data, considering diverse peptide characteristics like peptide classes, charge states, and sequence lengths. Our model attained a median cosine similarity score of 0.951 and 0.923 on the training and test data respectively. Contrary to existing deep learning models that often overlook a substantial part of peptide tandem mass spectra beyond the sequence b and y ion series, UniSpec can predict up to 75% of all measured fragment intensities (including unknown signals) in the raw experimental spectra. This represents a marked advancement from the 43.5% coverage achieved solely by b and y sequence ions in the NIST library spectra. For the evaluation of our model’s practical utility in predicting proteome-wide in-silico spectral libraries, we executed a benchmark test using a dataset of HeLa cells. UniSpec displayed a significant overlap of peptide identifications with the widely used search engine MS-GF+ and the NIST experimental spectral library, demonstrating its robust performance as a standalone peptide identification tool.

## Introduction

Determining peptide sequences and structures through tandem mass spectrometry relies on fragment ion analysis, which builds upon various fragmentation pathways derived from extensive theoretical and experimental research conducted by the mass spectrometry-based (MS) proteomics community^1-3^. Given the complexity and incomplete understanding of the multiple underlying mechanisms of peptide fragmentation, continued investigation is essential for deepening our comprehension of low-energy collision-induced peptide dissociation processes and furthering peptide identification in proteomics.

Leveraging the existing knowledge base, advancements in automated MS data analysis workflows have been made to accurately identify peptides^4-6^. Most sequence search engines use ad-hoc models to predict cleavage throughout all amide bonds on the peptide backbone, calculating masses for theoretical b and y ions. In general, these streamlined fragmentation algorithms have demonstrated great success in identifying peptides and proteins from shotgun proteomics analyses. Other tandem spectrum prediction models for peptides, such as MassAnalyzer^7^, simulate a variety of product ions found in most documented fragmentation pathways. Based on a mobile proton framework, this research sets the stage for innovative predictions of low-energy collision-induced dissociation in protonated peptides. Conventional machine learning approaches, such as PeptideArt^8^ and MS^2^PIP^9^, employ domain expertise-based algorithms for data parsing and have been developed to predict peptide fragment spectra utilizing rules derived from annotated experimental spectral data. Nevertheless, only a limited number of published predictive models are accessible in this area. Moreover, most existing bioinformatics tools are currently unable to identify various modified peptides, such as glycopeptides, phosphopeptides, and disulfide-bonded peptides, and interpret their product-ion spectra.

In contrast to the aforementioned rule-based models, deep learning (DL) techniques, a subset of machine learning that uses artificial neural networks to mimic the learning process of the human brain, have emerged as a more advanced solution for spectral prediction^10-11^. The recent advancements in this field^12-15^ have been fueled by modern shotgun proteomics generating large volumes of high-throughput peptide tandem mass spectrometry data, which provide ample data for training algorithms and architectures. The significant relationship between fragment spectral characteristics and peptide oligomeric structures presents an ideal context for the application of DL models. Deep learning outperforms more conventional machine learning algorithms due to its ability to learn intricate features and patterns from massive datasets without requiring feature engineering expertise^16^. Consequently, DL is particularly well-suited for the data-intensive and complex domain of MS proteomics. Fragment intensity prediction using DL has shown promise in advancing peptide identification with database search engines^17-18^ that match and score tandem mass spectra with possible peptides in proteomic samples, producing peptide spectrum matches (PSMs). By contrasting precise fragment intensity predictions with their corresponding PSMs from search engines, supplementary intensity-based scores can be determined, assisting in distinguishing true and false identifications. Similarly, DL has been employed to enhance peptide identification processes in library searches^19-20^, where query spectra are matched with previously identified library reference spectra. In such applications, DL models can create in-silico spectral libraries for any desired peptides, either as a substitute for or a supplement to experimental libraries.

In the literature, two methods of DL have been reported for predicting peptide spectra. The more common approach, represented by pDeep2^12^, Prosit^13^, AlphaPeptDeep^14^, and DeepDIA^19^, is the fragment intensity prediction method. These models predict fragment ion intensities by pre-defining 100 to 600 unique canonical ion categories, such as b2, y3, b2^2+^, y3^2+^, etc. These fragment ions are then converted into their theoretical *m/z* values for the specific peptide being predicted. These models require their training spectra to include annotations for the ions they will predict. In contrast to predicting intensities for predefined fragment ions, PredFull^15^ employs the binned method, predicting fixed width *m/z* bins with a spectral dispersion of 0.1 Da, outputting a total of 20000 intensities from *m/z* ranging from 0 to 2000. This method can be trained on any spectral data without requiring annotations, by transforming the high-precision *m/z* values acquired from modern high-resolution mass spectrometers into their corresponding low-precision bins of 0.1 Da. As individual bins often contain multiple high-precision fragment peaks with varying ion types and intensities, the collective intensities of the bins must be averaged or normalized to a single intensity value. While the binned method’s ion peaks can be assigned and up-resolved post hoc, ambiguities remain that complicate peptide identification applications. Furthermore, the predictions can also be negatively affected by contaminants and background ions.

The high mass accuracy of the fragment intensity prediction method is significantly more desirable than predicting low-resolution bins; however, predicted spectra containing only b and y ions fall short in identifying a broad range of peptides. Other ion types, such as a-ions, neutral losses, internal, and immonium ions, are also generated by low-energy collision-induced dissociation and are essential for interpreting intricate peptide spectra. Our goal is to develop a model that integrates extensive spectral coverage with high mass accuracy.

We present the development of a novel spectral prediction deep learning model (UniSpec) that predicts pre-defined ion fragments but expands the predicted ion output space far beyond canonical b and y ions; expanding to a size corresponding to the full fragmentation extent in experimental spectra. This model incorporates a basic permutation of ion types, peptide lengths, product charges, common neutral losses, and isotopic contributions, as well as immonium and internal ions. By leveraging high-quality NIST spectral libraries with extensive annotations of all detectable peptide fragment ions, the model’s output generates spectra with intensity predictions for 7919 isotope ions. Using UniSpec, we created an in-silico library of peptide tandem mass spectra covering the entire proteome to identify peptides in the HeLa dataset. Results from the comparison of the predicted library with sequence search engines and spectral libraries highlighted the promising capabilities of the model, suggesting its suitability as an effective alternative for analyzing proteomics data. Please email the authors directly if you wish to access the UniSpec source code and evaluate its datasets for the purposes of noncommercial research.

## Data and Methods

### Data source and preprocessing

#### Spectral libraries as data sources

We used NIST high-resolution mass spectral libraries from https://chemdata.nist.gov/dokuwiki/doku.php?id=peptidew:cdownload. These libraries were generated using raw data files acquired on the Orbitrap instrument models involving Velos, Elite, Q Exactive (QE), and Lumos, all publicly available on ProteomeXchange^21^. The NIST spectral library construction process is discussed elsewhere^22-23^. Briefly, when building the library, HCD spectra were searched using the MS-GF+^5^ with a precursor *m/z* tolerance of 20 ppm (i.e., parts per million). Identification results were collected with a false discovery rate of < 1%. Two types of library spectra are available: consensus and selected. The former is generated from the most frequent peaks in replicate spectra of a particular peptide with many different energies, while the latter contains the spectrum with the highest score selected from multiple identifications for a given peptide and energy. Consensus and selected spectra are indicated as “Consensus” and “Single”, respectively, by the first entry in the Comment section of the MSP file. These libraries are available in ASCII text format (MSP files) and are easy to use with many software programs.

#### Data preprocessing

We developed a set of metrics to evaluate the data quality of each mass spectral library, such as precursor *m/z* error, the ratio of unassigned abundance and noise peaks. These quality metrics are used to measure each spectrum, then the distributions and median**s** are calculated for each spectral library. Library-specific filters were determined based on the metrics’ median**s** and distribution**s** of each library. For example, for the human synthetic peptide library with a median of 0.9 ppm deviation of precursor *m/z* data and the human selected peptide library with a median of 2.1 ppm, the filter thresholds for precursors are conservatively set at 4.5 ppm and 14 ppm, respectively. The former was constructed from a high-quality single data source, while the latter was created using spectral data from many different laboratories and instruments over different time periods. These carefully determined filters were used to remove a fraction of the lowest-quality mass spectra.

Consistency between normalized collision energy (NCE) and the applied electron-volt (eV) is expected for the MS experiments using orbitrap instruments. The NCE setting, in the range of 10 - 100%, is used to determine the optimal eV for each of a wide range of masses during peptide fragmentation using a linear relationship between energy, mass, and charge (see Thermo Fisher Scientific Product Support Bulletin 104). To understand the relationship between NCE and eV, linear equations were derived by fitting the experimental data of the NIST reference materials^22,24^. We used these equations to investigate (1) the effect of NCE conversion on the optimal eV of the mass spectra, and (2) the reproducibility of peptide spectra across different instruments. The similarity between two spectra measured on different instruments was compared using the peptide spectral search software tool, NIST MS PepSearch (https://chemdata.nist.gov/dokuwiki/doku.php?id=peptidew:mspepsearch) to obtain the best-matched collision energy shifts, which were used as input for CE recalibration. Based on the analysis of more than 104,000 mirror spectra acquired on Lumos and QE, the highest percentage of best-matched spectra is found at an eV shift of six and an NCE shift of five between the two instruments. Similar comparisons were also conducted for other instruments, and the results show that Velos, Elite, and Lumos all require higher energy values to achieve similar levels of fragmentation efficiency as QE.

#### Dataset splitting

Utilizing data processing techniques, we compiled a cohesive spectral data collection tailored for model development, in which each raw mass spectrum was filtered, and energy recalibrated. Dataset splitting incorporated stratified and random sampling, along with sequence similarity discrimination, in a multi-step process. First, the included spectral libraries were divided into four subsets based on instruments; this allows us to draw more precise conclusions by ensuring that every instrument is properly represented in sampling. Next, two subsets were randomly chosen for validation and testing from each of the four subsets, with the remaining data designated for training. In the third step, sequences in the validation and test sets were compared to the training set and to each other using the Levenshtein edit distance algorithm^25^. Sequences from the two random subsets with normalized Levenshtein scores below 70, in the range of 0 (no match) to 100 (exact match), were considered sufficiently distinct from the training sequences, and thus are retained in the validation set (ValUniq) or test set (TestUniq). On the other hand, sequences with scores above 70 were either moved to the training set or to an additional test set (TestCommon) that mimics the training set. Finally, MS PepSearch was used to filter potentially highly similar spectra in validation and test sets to preserve data independence.

### High-resolution HCD fragmentation dictionary

In building a deep learning model that can predict the intensities of the full range of HCD fragment ion series, we developed a method for creating a high-resolution MS/MS fragment ion dictionary prior to training the model. The dictionary is derived from a comprehensive analysis of all fragment ions in the training data collected from the NIST peptide spectral reference library.

First, the training dataset is processed to extract a list of all measured/annotated neutral losses, internal, immonium ions, and amino acid fragment ions that result from cleavage of the peptide side chain and double backbone. Then, we generate all permutations of sequence ions, fragment sizes, charge states, peak isotopes, and a set of extracted neutral losses. Next, using the permutation list we enumerate the occurrence count of all candidate HCD MS/MS ions for the ion dictionary, across the entire training dataset. Leveraging peak frequency statistics, we can distill down the ion dictionary to only include ions that are sufficiently frequent in the training data. Ultimately, we utilize this refined dictionary to create a streamlined training dataset, which only incorporates the ions found in our dictionary for each spectrum.

When using the training set from this work, the permutation includes a, b, y, and precursor ion series with charges ranging from 1+ to 5+, up to 5 isotope peaks per ion, 23 extracted neutral losses, and up to 39-mer fragments. In addition, we added to the permutation list extracted internal fragment ions, which are defined by their starting position from the N-terminus, the total length of amino acids, and any neutral losses or modifications. The isotopic peaks and neutral losses were also considered for internal ions. In the low-mass region of the MS/MS spectrum, commonly observed immonium ions and miscellaneous fragments from individual amino acid residues were also added. This generated a list of a total of 79845 candidate isotope ion peaks. Of the over 89 million MS/MS fragment ions identified and annotated in the training data, we applied a conservative cutoff of 100 occurrences to the selections in the generated list, above which ions were added to the dictionary. This allows >99.9% of all MS/MS fragments to be included in model training. The resulting dictionary (see Supporting Information Tables S1a, b, and c for details) contains a total of 7919 isotopic peaks from a-ions (226), b-ions (2026), y-ions (3643), precursor ions (131), internal ions ion (1836) and iminium and residue fragment ion (57). All ion series except a-ions involve neutral losses.

### Model

This study uses an attention-based model, similar to the encoder structure of the transformer architecture^26^. The transformer was originally developed for natural language processing (NLP), which treats words or characters as separate tokens with features. Deep learning problems that work with amino acid sequences draw many parallels with NLP, including the inputs and tensor shapes that are operated on, and thus techniques from NLP can be transferred over to this problem.

The architecture of our model starts with an input tensor that embeds peptide sequence, modification, charge, and collision energy and ends with a one-dimensional output vector of prediction intensities for the 7919 isotope peaks in our dictionary. Sequence information is encoded as one-hot amino acid vectors along the channels dimension, across a maximum sequence length of 40 amino acids. A special token is added to the canonical 20 amino acids types as a null for absent amino acids short of the max length. The seven modifications used in the model (plus a null) also are represented as a one-hot vector. Charge and energy, which are peptide-level information, are replicated across the sequence dimension in order to embed them with the sequence and modification information. The charge is a one-hot vector up to charge eight, and the collision energy is a single float, in eV, divided by 100 to have a magnitude approximately 0 to 1. The input embedding is first projected to 256 channels and added to a learned positional embedding tensor. The resultant tensor is transformed through nine transformer-like blocks, each consisting of a self-attention block and a feed-forward block. The penultimate layer projects the 256 channels to 512 channels and the final layer projects to an output space of the length of our dictionary. Each predicted fragment intensity is generated by a sigmoid activation (between 0 and 1), and mean pooling over the sequence dimension. Throughout the network, we use batch normalization and various activation schemes to maintain stable training. Specific details on design and implementation can be found in Supporting Information Figure S1. The predicted intensities are then matched to a vector, also ordered by our ion dictionary indices, of the experimental spectrum’s corresponding ion intensities. The loss metric is to minimize the negative cosine similarity between the two vectors.

### Loss and evaluation metric

We calculate the model’s training loss using the negative cosine similarity between prediction vector X and target vector Y. The loss varies from -1 to 0, where -1 means X and Y are identical in peak’s *m/z* and intensity distribution. Zero signifies no shared peaks between the two spectra. We square root peak intensities before calculating cosine similarity, tempering the effect of very high intensity peaks. This preprocessing step improved spectral similarity analysis and is a common practice in MS/MS data analysis^27^.

In the validation process, we assess the similarity between two variable-length peak lists, such as those derived from MSP file representations of spectra, rather than using fixed-length vectors. This necessitates a similarity metric that is mostly annotation-agnostic, matching peaks only using their respective *m/z* values. We define the evaluation metric in Equation 1 as the cosine similarity score (CSS),

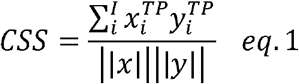

where *x*_*i*_ and *y*_*i*_ refer to predicted and experimental ion intensities, respectively. True positives *(TP)* refer to all the matched peaks between the two lists, also known as. Predicted peaks that do not match any experimental peaks are categorized as false positives (*FP*), and unmatched experimental peaks are labeled as false negatives (*FN*). In the denominator, “*x*” encompasses *TP* and *FP*, while “*y*” encompasses *TP* and *FN*. Peaks in the experimental spectrum that are unidentified are not included in the calculation, as their primary impact is to add noise and contamination.

The criteria for matching peaks are determined by product-ion *m/z* tolerances expressed in ppm. For instances under 800 *m/z*, peak pairs under 15 ppm are regarded as matches. Between 800 *m/z* and 1200 *m/z*, peak pairs under 20 ppm are seen as matches, and above 1200 *m/z*, those under 25 ppm are considered matches. When several predicted peaks match the reference peak within these tolerances, the predicted intensities are added up and the *m/z* closer to the reference is selected as the matching prediction. Predicted peaks that fall below the lowest *m/z* of the reference spectrum are not counted in the calculation. To mitigate the impact of some predominant precursor peaks on product ion similarity comparisons, they have been eliminated from all CSS calculations.

### Generation of the predicted tandem mass spectrum

To convert the predicted 7919-dimensional intensity vector to a tandem mass spectrum, data postprocessing includes calculating the theoretical *m/z* values of the isotopic peaks for all MS/MS ions and filtering. While the predicted vector contains the intensities of all ions in the dictionary, not all ions are obtainable for a given the peptide input. After including predictions above 0.1% maximum abundance, we filter based on plausibility, e.g., an n-mer peptide cannot generate any > (n-1)-mer fragments, fragment ions cannot have a higher charge than their precursors, etc. Furthermore, peptides do not yield modification-specific neutral losses in the absence of their corresponding modifications, e.g., the neutral loss of CH_3_SOH (64 Da) in precursor and/or product ions cannot occur on unoxidized Met-containing peptides. Also, we only included fragments in the 60 *m/z* to 1900 *m/z* range. After filtering, the remaining *m/z*, intensity, and annotations are written as predicted spectra in the MSP format, a text file used by the NIST mass spectral library.

### Data analysis and false discovery rate estimate

A dataset of human HeLa samples in PXD022287^28^ was downloaded from ProteomeXchange, which comprises peptides identified by MS-GF+^5^ from tryptic digests of HeLa cell lysates, along with their spectra acquired on an Orbitrap Fusion mass spectrometer. The raw data were searched with MS PepSearch against in-silico spectral libraries predicted by UniSpec with different collision energies as described above. For comparison purposes, a separate search against the NIST human peptide reference libraries was also conducted. Due to the numerous large in-silico spectral libraries and the limitation that the library search program can only simultaneously handle 12 MS/MS reference libraries, we searched separately the raw spectra against target and decoy spectral libraries instead of concatenating them together. This resulted in two sets of peptide spectrum matches (PSMs) returned, one from the target search and one from the decoy search.

To evaluate the false discovery rate (FDR) of our PSM results from separate searches, we employed the method suggested by Navarro et al.^29-30^ to compute the false positive proportion using the formula: (2xDB+DO)/(TB+TO+DB). In this equation, DB, DO, TB, and TO denote the number of PSMs obtained for decoy better, decoy only, target better, and target only, respectively, which are explained as follows. In the FDR assessment, when both target and decoy searches yielded identical PSMs and their scores exceeded a specific threshold, only one PSM was considered eligible based on their relative scores. If the decoy PSM score was higher than that of the target match, then it was classified as a false positive (DB or “decoy better”), while a target PSM was identified as a true positive (TB or “target better”) if its score was higher than the counterpart decoy match. In contrast, numerous PSMs with scores above the threshold were exclusive to either target matches (TO or “target only”) or decoy matches (DO or “decoy only”). After examining the raw data quality, we set the precursor tolerance at 5 ppm for all target and decoy PSMs. As a result, we determined a 1.0% FDR threshold at an MS PepSearch match score of 450.

## Results

### 1. Data strategy and deep learning datasets

The effectiveness of DL tools for spectral prediction depends on the quality of MS proteomics training datasets^31^. Our approach is to generate high-quality datasets based on the currently available NIST peptide spectral libraries created entirely from raw data stored in public data repositories (See Data and Methods). A total of more than 5 million HCD tandem mass spectra are included in these libraries with each spectrum uniquely defined by sequence, modification, charge, and collision energy. Libraries vary in how they are constructed, and the underlying data used; some incorporate chemical labeling experiments, others rely on label-free measurements, and they can be created using consensus or selected methods. NIST peptide libraries are comprehensive, curated mass spectral reference collections from various organisms and proteins useful for the rapid matching and identification of acquired MS/MS spectra. With such good resources for preparing AI datasets, we still need to address an important data preprocessing question: Can we directly combine all these different mass spectral libraries for model training? Since assembling training data in this way is a common practice, understanding this problem is especially important for developing reliable models.

To answer this question, we systematically examined two aspects of all relevant spectral libraries from a training perspective: (1) data quality and (2) applicability of different types of library spectra. First, we applied data quality metrics (See Data and Methods) to assess the overall quality of each of these libraries and used filters to remove spectra of low confidence based on their precursor *m/z* error, the ratio of unassigned fragment abundance, and noise peaks. Second, we trained the same model on different spectral libraries and compare their predictive performances. This comparison enabled us to assess the influence of various features of the training datasets on the overall model performance. Specifically, we evaluated the performance of the model on both consensus spectra (ModelCons) and selected spectra (ModelSel), separately, using the same validation set, and calculated the CSS for both models. In general, the predictions from ModelSel exhibited higher CSS values compared to ModelCons, indicating greater accuracy (See Supporting Information Figure S2). Our observations indicated that, besides the 1.7% of spectra that recorded identical CSS from both models, ModelSel surpassed ModelCons in terms of CSS in 60.6% of the cases, while ModelCons had higher CSS scores in 37.7% of the spectra. Since consensus spectra are the result of peaks occurring from many individual spectra with varying collision energies, we cannot use these spectra to train an accurate energy-sensitive model. For this reason, we determined that selected spectra were the most desirable for training.

Another equally important case is to study the effect of two different collision energy representations (NCE and eV) on model performance. The same model architecture was trained with NCE and eV embeddings separately and used to predict spectra from the validation set. Our results show that using eV instead of NCE yields 6 - 9% higher similarity scores across all charges and peptide lengths. This indicates that contrary to the popular use of NCE as model input^12-14,17^, eV is more sensitive to learning fragmentation patterns under a given energy. Further investigation of the inconsistency between NCE and eV in the MS datasets showed that in most HCD experiments when charges were misidentified during the automatic conversion from NCE to eV, about 5 - 20% of the mass spectra produce different spectra and eVs (see Supporting Information Figure S3). This explains why identical peptides with the same NCE could exhibit significantly different fragmentation patterns due to different applied eV energies.

The above comprehensive analysis resulted in selecting four “selected”-type, high-quality spectral libraries (e.g., including human, Chinese hamster ovary, human synthetic, and phosphorylated peptides) out of a total of eight label-free HCD spectral libraries available from the NIST website. To ensure that the model learns from true peptide identifications and clear fragmentation patterns, we filtered 13% of the spectra from these libraries based on large mass errors and a high noise peak ratio (see Data and Methods). We also adjusted the collision energy of 12% of the QE spectra by +5 eV and predicted 31% of the data with missing CE values. This approach improved the spectral data’s relevancy and consistency, resulting in a unified dataset of a total of 1.8 million relevant tandem mass spectra of around 0.8 million peptides suitable for training, validation, and testing. Seven common modifications were included (variable cysteine alkylation, methionine oxidation, N-terminal pyroglutamate from glutamine, N-terminal pyroglutamate from glutamate, N-terminal acetylation, pyro-carbamidomethyl N-terminal cysteine, and serine/threonine/tyrosine phosphorylation). The overall statistics of the datasets by the total number of spectra, peptide ions, and the percentage of modified peptide spectra are shown in Table 1. All datasets consist of tryptic peptides 6 to 40 amino acids long, charges 1+ to 8+, and seven common modifications. As is typical of proteomics data, a large majority of training spectra fall in a narrower range of characteristics than the extent of the data would suggest. For each dataset, more than 96% of the spectra involve charge states from 2+ to 4+, energies ranging from 15 - 70 eV, and peptide sizes from 6 to 30 residues.

**Table 1.** Dataset statistics.

| Datasets | # Spectra | # Peptide<br>ions | % Modifications |
| --- | --- | --- | --- |
| Training | 1715217 | 710109 | 30.3 |
| ValUniq | 12076 | 8468 | 45.3 |
| TestCommon | 49700 | 40479 | 23.8 |
| TestUniq | 6106 | 4987 | 46.8 |

Note: **spectra** =sequence+modification+charge+energy, **peptide ion** = sequence+modification+charge, ValUniq: validation set, TestUniq: test set, TestCommon: test set that has the same peptide subset used for training.

To answer the question posed at the top of this section, we argue that for an optimal model, the dataset quality should be more rigorous and targeted than simply including all spectra in currently available general spectral libraries. Spectral libraries have been designed to encompass all reliably identified peptide spectra. Nonetheless, given the intricate nature of the spectral analysis, there might be some low-quality mass spectra included that have inadequate peaks or considerable noise, even though they have been confirmed as true identifications. To train a deep learning model that can reliably learn peptide fragmentation patterns at specific energies, the dataset should first exclude false identifications as well as any low-quality spectra that may hinder correct pattern recognition. Furthermore, deep learning datasets should consist of raw experimental spectra acquired at specific energies rather than consensus spectra generated from replicates across multiple energies.

### 2. Implementation of the UniSpec attention model architecture

Figure 1 presents a schematic representation illustrating our strategy and implementation of our deep learning model. The six-step workflow begins with data preprocessing, where publicly available NIST libraries of peptide HCD MS/MS spectra are filtered and transformed into energy (eV) consistent format for input to the model. Next, the processed spectral data is split into training, validation, and test sets based on sequence dissimilarity. This division ensures a diverse range of sequence compositions for evaluation, helping to avoid overfitting and to improve the model’s ability to generalize to unseen peptide inputs. The third step involves generating a fragmentation ion dictionary based on the training data by surveying all annotated fragment ions in the raw dataset. In the fourth step, the training data is transformed into a streamlined data set, containing only the peaks from fragment ions present in our dictionary. This ensures that spectra can be read into target tensors as fast as possible during training. The fifth step entails building and implementing the model architecture on the training data, selecting appropriate hyperparameters, and optimizing the training process. Finally, the trained model was used to make predictions on the evaluation data, providing an estimate of its generalization ability. CSS was used as our evaluation score to quantify the model’s performance.

**Figure 1.**
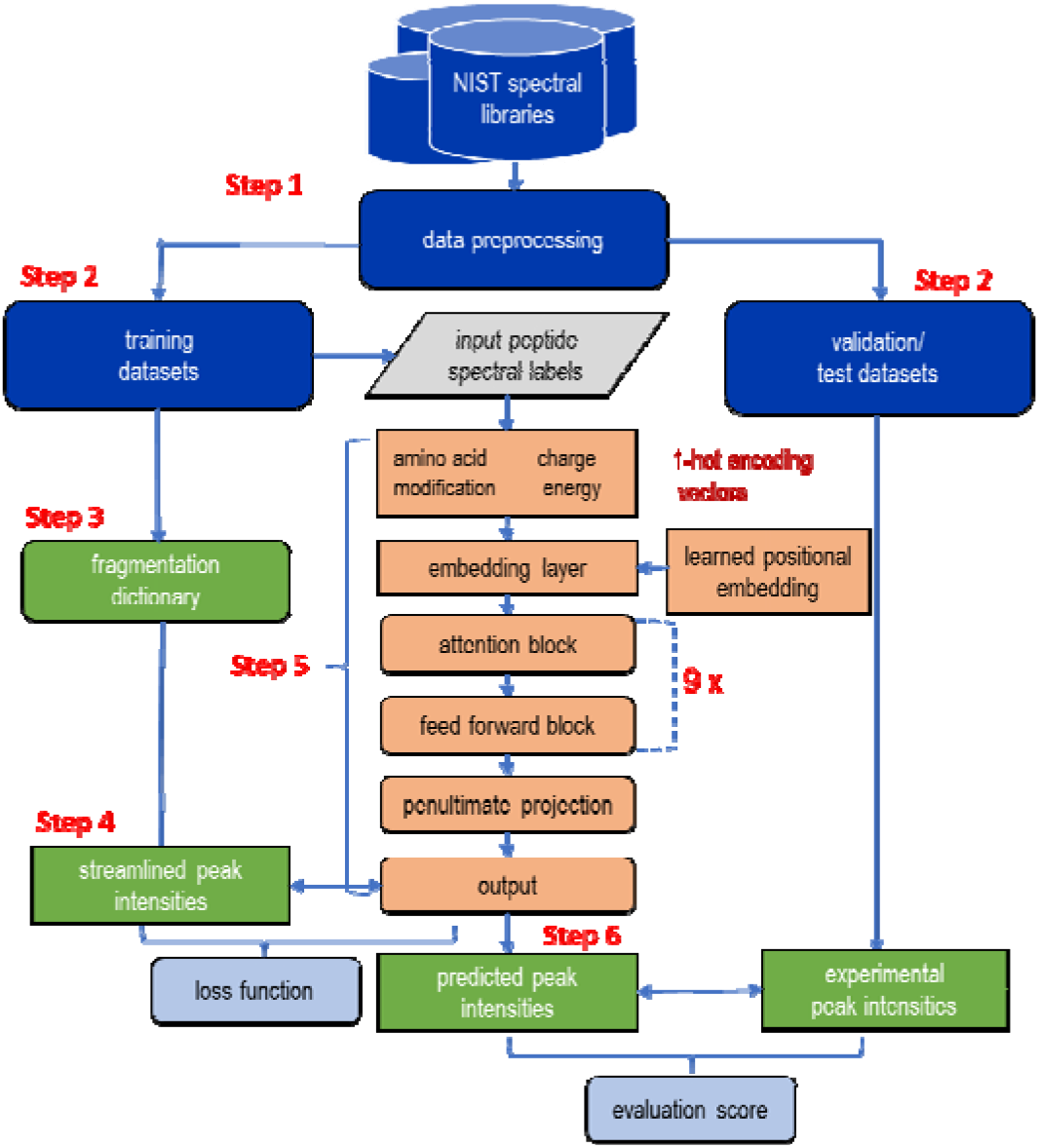
Implementation of the UniSpec attention model. The six-step workflow involves (1) data preprocessing, (2) splitting all data into training, validation, and testing datasets, (3) creating a fragmentation dictionary, (4) transforming data into streamlined peak lists, (5) building and training the model, and (6) predicting MS/MS spectra on the evaluation data (validation/test/external sets).

### 3. Evaluation of the model performance

#### 3.1 Evaluation by peptide characteristics

To evaluate the accuracy of the model’s predictions, we compared its performance on testing data with that of the training data. We predicted spectra from TestCommon (i.e., closely resembling the training data) and TestUniq (i.e., dissimilar from the training data) and calculated their CSS. Figure 2a shows that the overall predictions on the training data obtain a median CSS of 0.951, while the predictions on the test data yield a median CSS of 0.923, which is within 0.03 points of the training accuracy. This small disparity between training and testing CSS assuages concerns of overfitting and illustrates that UniSpec is a good general predictor of fragment ion spectra.

**Figure 2a.**
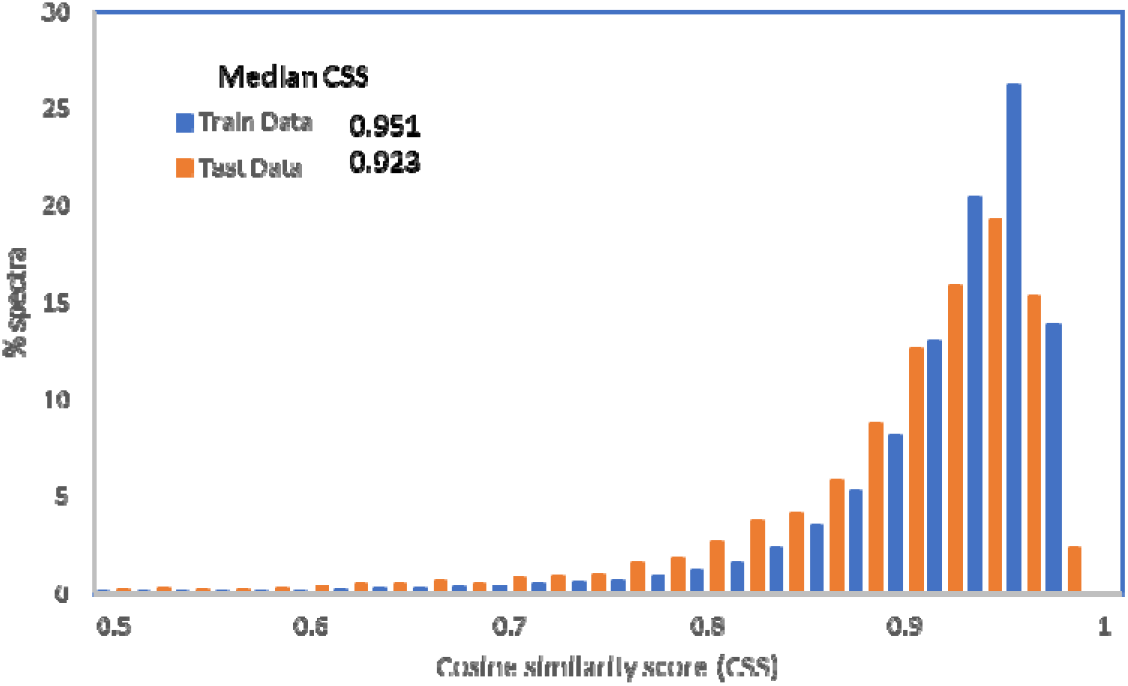
Comparing performance on testing data to training data.

We also extended this comparative investigation of model performance to training and test data across various peptide characteristics, such as peptide classes, charge states, and sequence lengths (Table 2 and Figures 2b and 2c). All comparisons are within 0.03 between testing and training accuracy, except 0.05 for high charge 5+ and 0.04 for peptide sizes of 36 to 40 amino acid residues. Table 2 provides the CSS scores for the five major peptide classes produced by trypsin digestion^22,32^. Class 1 and Class 2 are products of simple tryptic cleavage with or without common modifications and therefore are expected to represent the bulk of the ion intensity in tryptic digests of proteins. Class 3 and 4 peptides are usually produced by incomplete digestion. Class 5 peptides have one terminus produced by non-tryptic cleavage. The last three classes were less abundant than the first two, although more peptides were commonly observed in LC-MS/MS analyses. Unlike the first two classes, Classes 3 and 4 include one or more missed cleavage sites and have therefore longer and more complex sequences, resulting in intricate fragmentation patterns. Class 5 consists of irregularly cleaved peptides, which are less sought and studied in LC-MS studies. In general, modified peptides add more complexity to the structure and fragmentation than unmodified peptides. As expected, unmodified simple tryptic peptides (Class 1) scored the highest in the training and test sets. Compared to Class 1’s performance, other classes obtain reduced scores consistent with the diversity of their sequences and fragments. Overall, all score differences between Class 1 and the others are within 0.04 for the training and test data, showing that the model can handle a wide variety of different peptides.

**Figure 2b.**
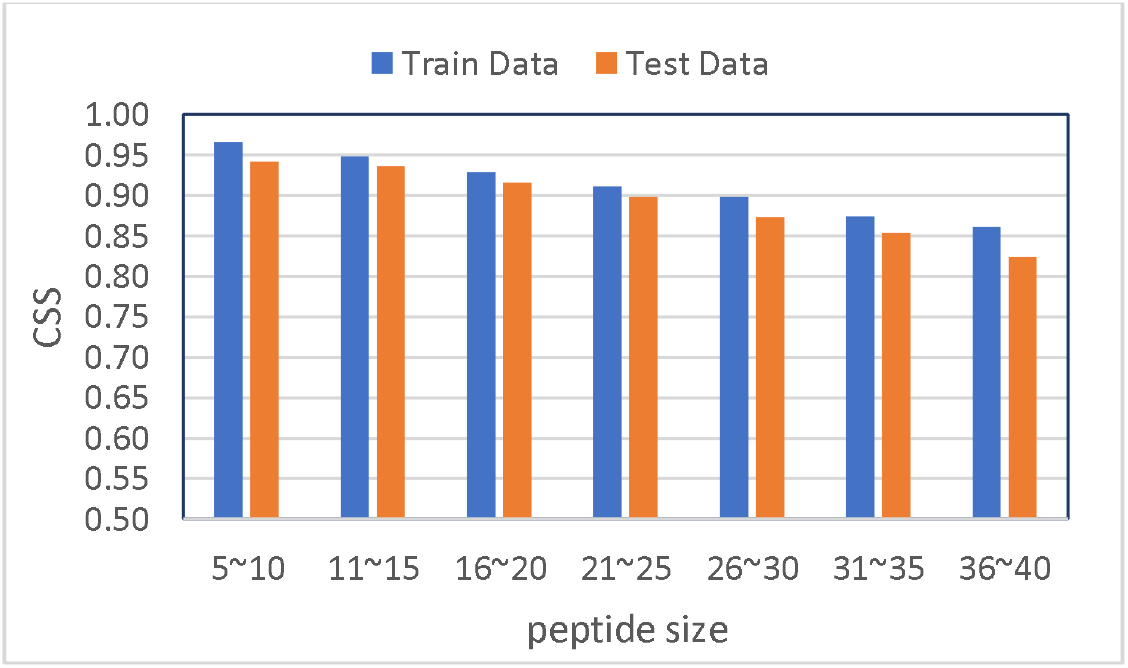
CSS distribution over peptide sizes from 5 to 40.

**Figure 2c.**
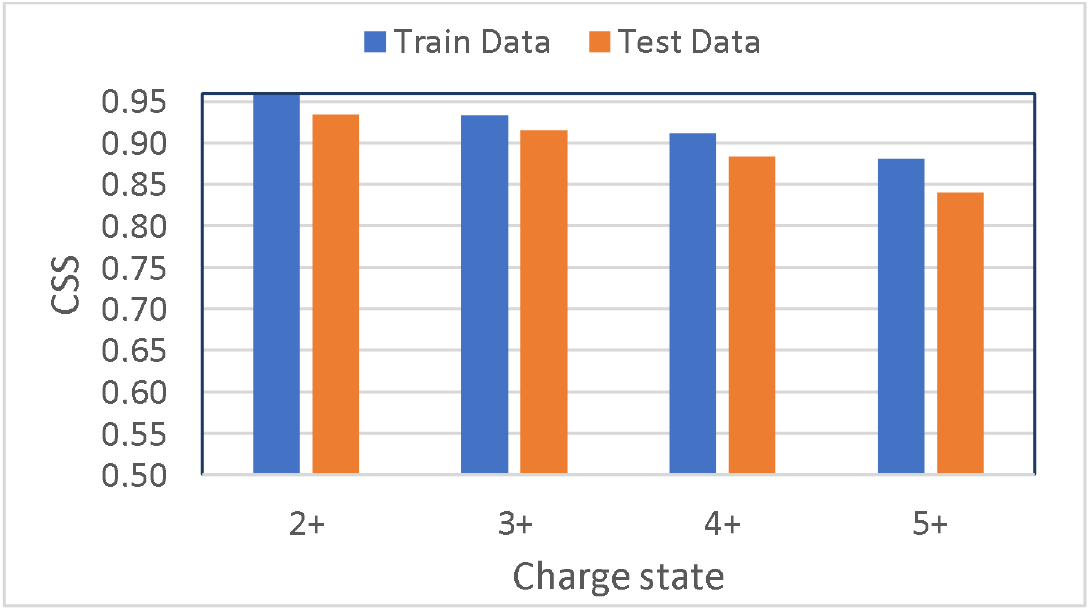
CSS distribution over charge states from 2+ to 5+.

**Table 2.** The median cosine similarity score was calculated for major peptide groups in TestCommon (most similar to the training data) and TestUniq (dissimilar to the training data).

| Group | Description | TestCommon | TestUniq |
| --- | --- | --- | --- |
| 1 | Tryptic_unmodified | 0.964 | 0.947 |
| 2 | Tryptic_modified | 0.942 | 0.933 |
| 3 | Miscleaved_unmodified | 0.936 | 0.919 |
| 4 | Miscleaved_modified | 0.924 | 0.905 |
| 5 | Semitryptic | 0.925 | 0.918 |

This work was in part inspired by Prosit^15^, a popular deep-learning model that has been effectively applied in proteomics studies. Based on the similar evaluation method used by both models (for each model’s respective predefined ion series and testing datasets), our model’s testing accuracy is comparable to the median angular similarity from the original Prosit RNN’s 0.908^13^ and from a recent Prosit Transformer’s 0.929^33^.

#### 3.2 Evaluation by overall peptide HCD fragmentation

To date, deep learning models have largely excluded a substantial fraction of HCD fragment ions beyond the typical sequence b and y ion series, mainly due to the challenges of understanding and annotating complex fragment ion types under collision-induced dissociation. However, these additional ions generated by alternative fragmentation mechanisms, namely neutral losses, internal ions, immonium ions, and side chain fragment ions, have been widely observed^34^. The fragmentation statistics obtained from the NIST spectral library revealed that besides the primary b/y ions, the additional ions collectively exhibit an average abundance of 28% relative to the total spectrum intensity in the 824529 spectra of the major unmodified peptides. Moreover, for biologically modified peptides this percentage increases to 32% across 124723 spectra for oxidized peptides and 37% across 66918 spectra for phosphopeptides. The detailed contributions can be found in Supporting Information Figure S4. These data suggest that the intricate HCD spectral content may encompass substantial contributions from various fragment ion types, extending beyond the range of the b/y ion series.

Predictions based on the validation/test sets demonstrated that UniSpec is capable of predicting various fragment types such as (1) sequence a/b/y ions, (2) neutral losses, (3) internal fragments, (4) immonium ions and side chain fragments, and (5) precursor and related ions. The contribution of each of these ion types to the overall intensity coverage is illustrated in Figure 3, showing that the relative abundances of the five ion types mentioned above are 43.5%, 13.6%, 3.9%, 3.8%, and 10.7%, respectively. The remaining 24.5% intensity in the figure is from observed unknown ions, which collectively constitute potentially unannotated, contaminant, and noise peaks. Based on our analysis, by aiming to predict all HCD fragment ion types, UniSpec achieves an average predicted coverage of 75.5% of all measured fragment abundances (including unknown signals), a significant increase compared to the 43.5% total coverage by sequence ions only.

**Figure 3.**
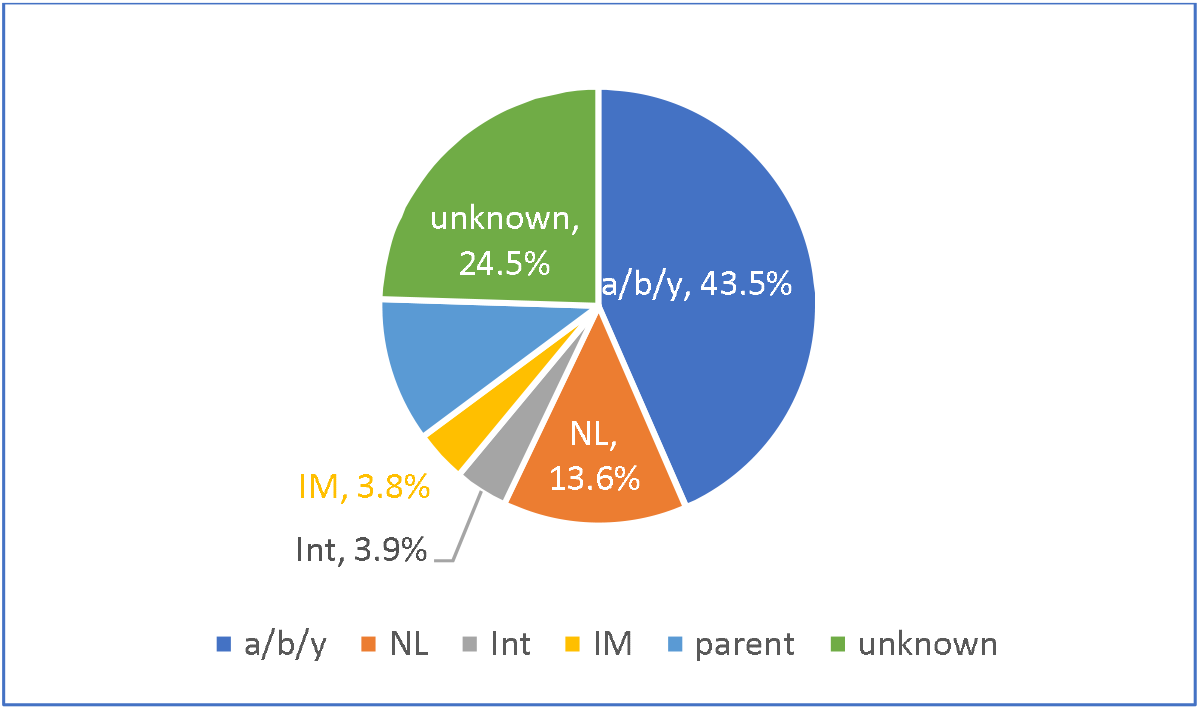
Predicted intensity coverage of the five major ion types in HCD spectra in the validation set of 12076 spectra. Note: Five ion types are: (1) sequence ions (a/b/y), (2) neutral losses (NL), (3) Internal fragme ts (Int), (4) immonium ions and side chain fragments (IM), and (5) precursor and related ions (Parent). The remaining unknown peaks in each validation spectrum include potentially unannotated, contaminant, and noise signals.

Figure 4 shows two examples of how our model predicts various fragment ion series for unmodified and covalently modified peptides. Figure 4a shows a head-tail plot comparing the predicted spectrum (bottom) of the triply-charged unmodified peptide ion SSSHTSLFSGSSSSTK with the corresponding experimental spectrum (top). The computed spectral similarity score was 0.923, which is representative of the median predictive performance over the entire test set of peptides. Overall, this is a relatively complex MS/MS prediction; the model accurately predicts all major b and y ion series, which account for 50% of the total experimental fragment abundance. Significant alternative fragments were also well-predicted, such as the neutral losses tied to numerous serine and threonine residues. These predictions contributed to a substantial H2O loss (29% of total abundance) in both the precursor and b and y series. In addition, relatively intense a-series peaks (5% of total abundance), such as *m/z* 336.6 and 410.2, and the frequently observed phenylalanine immonium ion at *m/z* 120 are predicted. As shown in Figure 4a, except for a few peaks below 3% abundance, all observed fragments, including isotopic peaks, were correctly predicted, with highly similar fragment intensities between predictions and measurements.

**Figure 4a.**
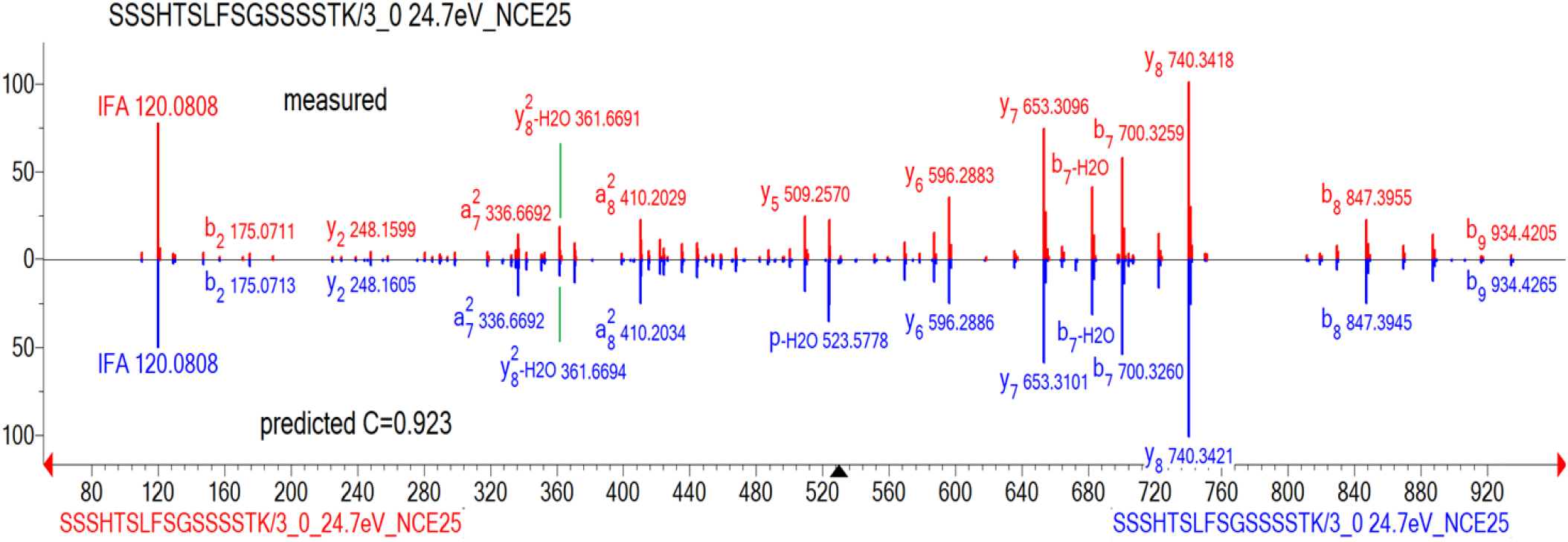
Head-to-tail plot showing a predicted peptide HCD spectrum (bottom, blue) of a triply charged ion of SSSHTSLFSGSSSSTK at 24.7eV/NCE25, and the matching experimental spectrum (top, red).

In Figure 4b we show an example of a covalently modified peptide, also at median predictive CSS of the test set. UniSpec can predict the two highly selective cleavages of the N-terminal side to proline residues, resulting in the two most dominant fragments y_5_ at *m/z* 588.292, and the neutral loss of CH_3_SOH from y_10_ ions. For this doubly charged peptide ion with an oxidized methionine, the model has learned to generate (under low proton mobility in this case) the dominant neutral loss of methane sulfenic acids (CH_3_SOH, 64 Da) from the side chain of methionine sulfoxide residues. This was demonstrated by the prediction of multiple large, consecutive peaks in the higher mass region, corresponding to the loss of CH_3_SOH from the y_9_ to y_12_ ion series.

**Figure 4b.**
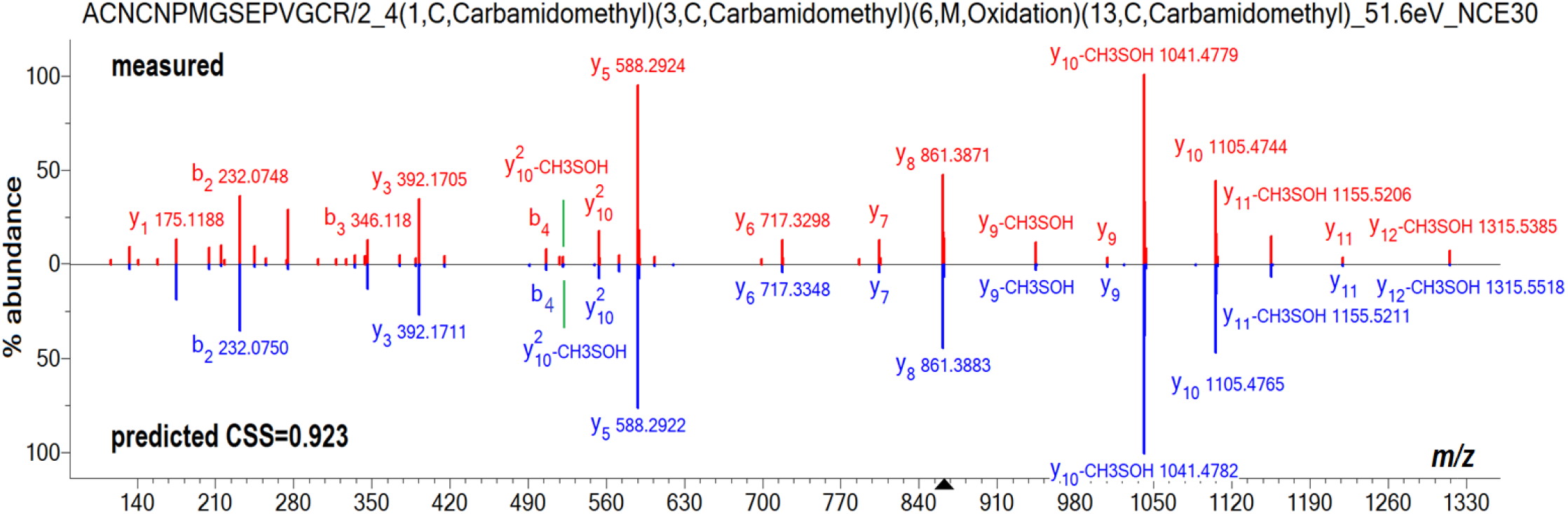
Head-to-tail plot showing a predicted peptide HCD spectrum (bottom, blue) of a doubly charged peptide ACNCNPMGSEPVGCR with three Carbamidomethyl cysteine sites and oxidized methionine at 51.6 eV/NCE30, and the matching experimental spectrum (top, red).

#### 3.3 Evaluation by investigating low-score predictions

In the previous sections, we discussed the advantages of UniSpec. In this section, We comprehend the weaker aspects of model performance by carefully examining the predictions with lower CSS produced by the validation and test datasets. Of the 18166 predicted spectra, we identified 371 of the lowest scores when using a cut-off score of 0.55 (2% of the total spectra). Detailed analysis of these low-scoring spectra revealed that while the model correctly predicted many fragment ions, their intensities varied widely from the experimental spectrum. One category of inaccurate predictions involves spectra that are dominated by parent peaks because of little to no dissociation of the precursor ion under the given collision energies (See Supporting Information Table S2 and Figure S5). In these cases, most of the detected fragment ions are minor or trace ions. Such peptides typically have one or more internal basic residues and prolines in their sequence, and the applied collision energies are below the threshold needed to mobilize a proton for charge-directed cleavage. These experimental spectra are often of low quality, presenting challenges for model training and peptide identification. Another difficult case in prediction was from spectra representing 8 - 13% of large peptides of 35 or more amino acids. Such spectra may be characterized by complex fragmentation patterns within the peptide backbone and side chains (See Supporting Information Figure S6). Note that there may be cases where the CSS is low because a low-quality experimental spectrum deviated greatly from the correct prediction.

### 4. Benchmarking proteome-scale in-silico spectral libraries on a HeLa dataset

Experimental spectral libraries can significantly contribute to the precise detection and quantification of unmodified/modified peptides in targeted quantitative proteomics, data-independent acquisition (DIA), and post-translational modifications. Given their effectiveness, we embarked on generating in-silico spectral libraries, employing a series of procedures outlined in Supporting Information Document S1. The aim was to evaluate the feasibility of using in-silico library searches as a viable alternative for identifying peptides from MS/MS spectra in real-world proteomics experiments. Specifically, we assessed the performance of a UniSpec generated in-silico library utilizing a human HeLa dataset (PXD022287), reported by Zeng and Ma^28^. The authors identified a large set of HeLa peptides from tryptic digests of HeLa cell lysates utilizing MS-GF+ (see Method). We compared the results of our in-silico library for peptide identification to other popular data analysis software tools, such as MS-GF+ (i.e., sequence search) and the MS PepSearch/NIST Human Library (i.e., spectral search).

We analyzed the raw data using MSPepSearch against the in-silico spectral library, predicted by UniSpec using protein sequence databases (refer to Methods), and, separately, the NIST consensus human peptide library, created from 10000 MS/MS raw data files acquired on Orbitrap mass spectrometers^23^. To ensure a fair comparison, we implemented similar search parameters as in the initial analysis, filtering the predicted and experimental spectral library results with a precursor *m/z* error of 5 ppm and a fixed cysteine carboxymethylation modification. Through this process, we identified 33745 and 33387 distinct peptides from the predicted and experimental libraries, respectively, both with a match score above 450 (out of 1000) and a 1% FDR. We compared these results with the 34659 peptides obtained from MS-GF+ with a Q-value cutoff of <0.01, as stated by the authors^28^.

The comparison results in Figure 5 demonstrate that the in-silico library exhibits a strong standalone performance on this dataset, yielding 30468 (87.9%) and 32904 (98.6%) significant identification overlaps with MS-GF+ and the NIST human peptide library, respectively. Despite this, the outcomes of the three search methods display noticeable differences, which are further analyzed and discussed in Supporting Information Figure S7. Briefly, the predicted spectral library identified 9.5% of the different peptides not detected by MS-GF+ and 2.5% of the unique peptides missed by the NIST spectral library. On the other hand, our library could not identify 12.4% and 1.4% of the peptides that MS-GF+ and the NIST spectral library, respectively, had detected. It is worth noting that these results not only mirror the quality and coverage of spectral libraries but also the performance of the spectral search engine used, which is not the focus of this study. Overall, these findings suggest that the in-silico library search is on par with MS-GF+ and the NIST human library, underscoring its potential as a supplementary tool in conjunction with traditional database and experimental spectral library searches.

**Figure 5.**
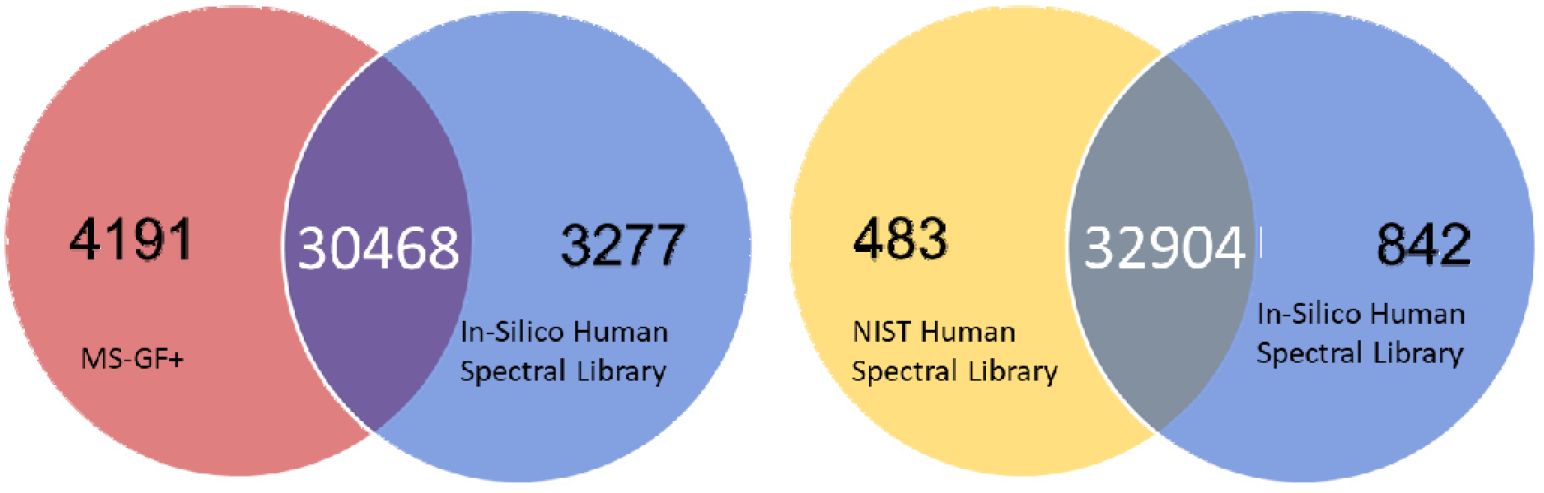
illustrates a performance comparison of predicted libraries, MS-GF+ database search, and NIST Human peptide libraries on the HeLa dataset. The same peptides identified by separate in-silico and NIST library searches amount to 30,468, while separate in-silico and MS-GF+ database searches reveal 32,904 shared peptides. The figures highlighted in black indicate the unique peptides identified exclusively by each respective method.

## Conclusion

Our study established a sophisticated deep learning neural network that reliably predicts the intensities of numerous, complex HCD MS/MS fragment ion types, surpassing the capabilities of existing machine learning tools that limit their focus to the b/y ion series. Our goal was to employ such a comprehensive model in the creation of in-silico spectral libraries to facilitate the identification of peptides from complex mass spectrometric data.

To achieve this goal, our extensive data preprocessing steps combined with the training of deep learning models help identify the key elements required to successfully train energy-sensitive models. These are: (1) choosing the selected best spectra for training, rather than consensus spectra constructed from average collision energies; (2) using eV embeddings to replace the partially erroneous NCEs observed in LC-MS/MS experiments. This allowed us to compile high-quality training, validation, and testing datasets from a diverse range of real-world peptides from the curated NIST mass spectral libraries. Based on inputs of sequence, charge, and collision energy (eV), as well as comprehensive peak annotations in the training data, UniSpec efficiently learns consistent intensity patterns for various product-ion types as ground truth, resulting in reliable peptide spectrum predictions. We evaluated UniSpec through a test set and tested its application on a HeLa peptide case study, which revealed its ability to predict all known fragment ion types. These included sequence a/b/y ions, neutral losses, internal fragments, immonium ions, amino acid side chain fragments, and precursor ions. This method shows a significant improvement over current machine learning tools, elevating the average coverage of all measured known/unknown signal intensities from 43.5% (with sequence ions only) to 75%. This technique ultimately extends the usability of current predictive models to a broad range of peptides and modified peptides engaged in intricate proteomics studies.

While existing deep learning models have been shown to enhance peptide search engines by rescoring peptide spectrum matches, we believe their primary applications will involve in-silico spectral libraries applied to the workflows of quantitative proteomics, data-independent acquisition, post-translational modification analysis, and more. We tested our proteome-wide in-silico spectral library on a HeLa dataset and showed that, in addition to enabling broad applicability by constructing the library for the entire proteome, our library building can be optimized according to the instrument and collision energies from the data that needs to be analyzed. This shows the flexibility and specificity advantages of predicted spectral libraries over building experimental spectral libraries. The results suggested that an UniSpec-predicted library is a viable alternative to a sequence database and an experimental spectral library. The possibilities of predicted spectral libraries are highly promising, given their strong performance in peptide identifications, energy optimization capability, and easily customizable potential for proteomics studies of interest.

## Associated Content

### Supporting Information

Model architecture schematic (Figure S1); Evaluation of two models of ModelSel and ModelCons using the same validation set (Figure S2); Conversion errors from NCE to eV observed in the NIST antibody NCE 24 experiment (Figure S3); Average abundance coverage of the three major ion types in unmodified peptide spectra (Figure S4 a); Average abundance coverage of the three major ion types in oxidized peptide spectra (Figure S4 b); Average abundance coverage of the three major ion types in phosphopeptide spectra (Figure S4 c); The head-to-tail plots of the predicted and experimental spectrum from the test set (Figure S5); The head-to-tail plot of the predicted and experimental spectrum from the validation set (Figure S6); Comparison of predicted and experimental spectra of a triply-charged peptide ion (Figure S6); and Generation of in-silico spectral libraries (Document S1) (PDF)

Content of High-resolution HCD fragmentation dictionary (Table S1 a); Neutral losses are used in the dictionary (Table S1 b); Commonly observed immonium ions and miscellaneous fragments relative to individual amino acid residues (Table S1 c) Examples of inaccurate predictions with dominant parent peaks (Table S2) (XLSX)

## Acknowledgments

The authors would like to thank William Wallace, Sergey Sheetlin, and Meghan Burke Harris for their support with data collection and interesting discussions. We would also like to thank Karl Irikura and Trina Mouchahoir for their detailed comments on the manuscript. This work was supported solely with NIST funds. Certain commercial instruments or software are identified in this paper to specify the experimental procedure and data analysis workflow adequately. Such identification is not intended to imply recommendation or endorsement by the National Institute of Standards and Technology, nor is it intended to imply that the materials or instruments identified are necessarily the best available for the purpose.

## References

1. Wysocki, V. H.; Tsaprailis, G.; Smith, L. L.; Breci, L. A. Mobile and Localized Protons: A Framework for Understanding Peptide Dissociation. J. Mass Spectrom. 2000, 35, 1399–1406.

2. McLafferty, F. W.; Turec?ek, F. Interpretation of Mass Spectra 4th ed.; 1993, University Science Books: Mill Valley, CA.

3. Kapp, E. A.; Schu1tz, F.; Reid, G. E.; Eddes, J. S.; Moritz, R. L.; O’Hair, R. A. J.; Speed, T. P.; Simpson, R. J. Mining a Tandem Mass Spectrometry Database to Determine the Trends and Global Factors Influencing Peptide Fragmentation. Anal. Chem. 2003, 75, 6251–6254.

4. Perkins, D. N.; Pappin, D. J.; Creasy, D. M.; & Cottrell, J. S. Probability-based protein identification by searching sequence databases using mass spectrometry data. Electrophoresis, 1999, 20 (18), 3551–3567.

5. Kim, S.; Pevzner, P. A. MS-GF+ makes progress towards a universal database search tool for proteomics. Nat Commun. 2014, 5, 5277.

6. Cox, J.; and Matthias, Mann. MaxQuant enables high peptide identification rates, individualized p.p.b.range mass accuracies and proteome-wide protein quantification. Nature biotechnology. 2008, 26 (12), 1367–72.

7. Zhang, Z. Prediction of low-energy collision-induced dissociation spectra of peptides. Anal Chem. 2004, 76 (14), 3908–3922.

8. Arnold, R. J.; Jayasankar, N.; Aggarwal, D.; Tang, H.; Radivojac, P. A machine learning approach to predicting peptide fragmentation spectra. Pac Symp Biocomput. 2006, 219–230.

9. Degroeve, S.; and Martens, L. MS^2^PIP: a tool for MS/MS peak intensity prediction. Bioinformatics. 2013, 29, 3199–3203

10. Cox, J. Prediction of peptide mass spectral libraries with machine learning. Nat Biotechnol. 2023, 41 (1), 33–43.

11. Wen, B.; Zeng, W. F.; Liao, Y.; Shi, Z.; Savage, S. R.; Jiang, W.; Zhang, B. Deep Learning in Proteomics. Proteomics. 2020, 20, 21–22.

12. Zeng, W. F.; Zhou, X. X.; Zhou, W. J.; Chi, H.; Zhan, J.; and He, S. M. MS/MS Spectrum Prediction for Modified Peptides Using pDeep2 Trained by Transfer Learning. Anal Chem., 2019, 91(15), 9724–9731.

13. Gessulat, S.; Schmidt, T.; Zolg, D.P.; Samaras, P.; Schnatbaum, K.; Zerweck, J.; Knaute, T.; Rechenberger, J.; Delanghe, B.; Huhmer, A.; Reimer, U.; Ehrlich, H.C.; Aiche, S.; Kuster, B.; Wilhelm, M. Prosit: proteome-wide prediction of peptide tandem mass spectra by deep learning. Nat Methods. 2019,16 (6), 509–518.

14. Zeng, W. F.; Zhou, X. X.; Willems, S.; Ammar, C.; Wahle, M.; Bludau, I.; Voytik, E.; Strauss, M. T.; and Mann, M. AlphaPeptDeep: a modular deep learning framework to predict peptide properties for proteomics. Nat. Commun. 2022, 13 (1), 7238.

15. Liu, K.; Li, S.; Wang, L.; Ye, Y.; Tang, H. Full-Spectrum Prediction of Peptides Tandem Mass Spectra using Deep Neural Network. Anal Chem. 2020, 92 (6), 4275–4283.

16. Shinde, P. P.; & Shah, S. A review of machine learning and deep learning applications. In 2018 Fourth international conference on computing communication control and automation (ICCUBEA) 2018, (pp. 1–6). IEEE.

17. Zolg, D. P.; Gessulat, S.; Paschke, C.; Graber, M.; Rathke-Kuhnert, M.; Seefried, F.; et al. INFERYS rescoring: boosting peptide identifications and scoring confidence of database search results. Rapid Commun. Mass Spectrom. 2021, e9128.

18. Wilhelm, M.; Zolg, D. P.; Graber, M.; Gessulat, S.; Schmidt, T.; Schnatbaum, K.; Schwencke-Westphal, C.; Seifert, P.; de Andrade Krätzig, N.; Zerweck, J.; Knaute, T.; Bräunlein, E;, Samaras, P.; Lautenbacher, L.; Klaeger, S.; Wenschuh, H.; Rad, R.; Delanghe, B.; Huhmer, A.; Carr, S. A.; Kuster, B. Deep learning boosts sensitivity of mass spectrometry-based immunopeptidomics. Nature communications. 2021, 12 (1), 3346.

19. Yang, Y.; Liu, X.; Shen, C.; Lin, Y.; Yang, P.; Qiao, L. In-silico spectral libraries by deep learning facilitate data-independent acquisition proteomics. Nature communications, 2020, 11 (1), 146.

20. Sinitcyn, P.; Hamzeiy, H.; Salinas Soto, F.; Itzhak, D.; McCarthy, F.; Wichmann, C.; Steger, M.; Ohmayer, U.; Distler, U.; Kaspar-Schoenefeld, S.; Prianichnikov, N.; Yilmaz, S.; Rudolph, J. D.; Tenzer, S.; Perez-Riverol, Y.; Nagaraj, N.; Humphrey, S. J.; Cox, J. MaxDIA enables library-based and library-free data-independent acquisition proteomics. Nature biotechnology, 2021, 39 (12), 1563–1573.

21. Vizcaíno, J.; Deutsch, E.; Wang, R. et al. ProteomeXchange provides globally coordinated proteomics data submission and dissemination. Nat Biotechnol 2014, 32, 223–226.

22. Dong, Q.; Liang, Y.; Yan, X.; Markey, S. P.; Mirokhin, Y. A.; Tchekhovskoi, D. V.; Bukhari, T. H.; Stein, S. E. The NISTmAb tryptic peptide spectral library for monoclonal antibody characterization. mAbs 2018, 10, 354–369.

23. Sheetlin, S. L.; Wang, G.; Tchekhovskoi, D. V.; Zhang, Z.; Stein, S. E. Filtering and optimization of peptide tandem mass spectral libraries. The ASMS 2020 conference proceedings, 2020, June.

24. Simón-Manso, Y.; Lowenthal, M. S.; Kilpatrick, L. E.; Sampson, M. L.; Telu, K. H.; Rudnick, P. A.; Mallard, W. G.; Bearden, D. W.; Schock, T. B.; Tchekhovskoi, D. V.; Blonder, N.; Yan, X.; Liang, Y.; Zheng, Y.; Wallace, W. E.; Neta, P.; Phinney, K. W.; Remaley, A. T.; Stein, S. E. Metabolite profiling of a NIST Standard Reference Material for human plasma (SRM 1950): GC-MS, LC-MS, NMR, and clinical laboratory analyses, libraries, and web-based resources. Anal. Chem. 2013, 85, 11725–11731.

25. Navarro, G. A. guided tour to approximate string matching. ACM Computing Surveys 2001, 33 (1), 31–88.

26. Vaswani, A.; Shazeer, N.; Parmar, N.; Uszkoreit, J.; Jones, L.; Gomez, A.N.; Kaiser, L.; Polosukhin, I. Attention is all you need. Adv Neural Inf Process Syst. 2017, 5998–-6008.

27. Stein, S.E.; Scott, D.R. Optimization and testing of mass spectral library search algorithms for compound identification, J. Am. Soc. Spectrom. 1994, 5 (9), 859–866

28. Zeng, X.; & Ma, B. MSTracer: A Machine Learning Software Tool for Peptide Feature Detection from Liquid Chromatography-Mass Spectrometry Data. Journal of proteome research, 2021 20 (7), 3455–3462.

29. Jung, K. Statistical methods for proteomics. Methods Mol Biol. 2010, 620, 497–507.

30. Navarro, P.; Vázquez, J.; A refined method to calculate false discovery rates for peptide identification using decoy databases. J Proteome Res. 2009, 8 (4), 1792–6.

31. Palmblad, M.; Böcker, S.; Degroeve, S.; Kohlbacher, O.; Köll, L.; Noble, W. S.; & Wilhelm, M. Interpretation of the DOME Recommendations for Machine Learning in Proteomics and Metabolomics. Journal of proteome research, 2020, 21 (4), 1204–1207.

32. Dong, Q.; Yan, X.; Kilpatrick, L.E.; Liang, Y.; Mirokhin, Y.A.; Roth, J.S.; Rudnick, P.A.; Stein, S.E. Tandem mass spectral libraries of peptides in digests of individual proteins: Human Serum Albumin (HSA). Mol. Cell. Proteomics. 2014, 13 (9), 2435.

33. Ekvall, M.; Truong, P.; Gabriel, W.; Wilhelm, M.; Käll,L. Prosit Transformer: A transformer for Prediction of MS2 Spectrum Intensities. J Proteome Res 2022, 21 (5), 1359–1364.

34. Michalski, A.; Neuhauser, N.; Cox, J.; Mann, M. A systematic investigation into the nature of tryptic HCD spectra. J Proteome Res. 2012, 11 (11), 5479–91.

35. Shao, W.; & Lam, H. Tandem mass spectral libraries of peptides and their roles in proteomics research. Mass spectrometry reviews, 2017, 36 (5), 634–648.

36. Ammar, C.; Berchtold, E.; Csaba, G.; Schmidt, A.; Imhof, A.; & Zimmer, R. Multi-Reference Spectral Library Yields Almost Complete Coverage of Heterogeneous LC-MS/MS Data Sets. Journal of proteome research, 2019 18 (4), 1553–1566.

37. Heil, L. R.; Fondrie, W. E.; McGann, C. D.; Federation, A. J.; Noble, W. S.; MacCoss, M. J.; & Keich, U. Building Spectral Libraries from Narrow-Window Data-Independent Acquisition Mass Spectrometry Data. Journal of proteome research, 2022, 21 (6), 1382–1391.

38. Yang, Y.; Liu, X.; Shen, C.; Lin, Y.; Yang, P.; & Qiao, L. In-silico spectral libraries by deep learning facilitate data-independent acquisition proteomics. Nature Communications, 2020, 111), 146.

