## Supplementary Figures for "UniSpec: A Deep Learning Approach for Predicting Energy-Sensitive Peptide Tandem Mass Spectra and Generating Proteomics-Wide In-Silico Spectral Libraries"

Table of Contents

Supporting Figures

Figure S1 – Model architecture schematic

Figure S2 – Evaluation of two models of ModelSel and ModelCons using the same validation set

Figure S3 – Conversion errors from NCE to eV observed in the NIST antibody NCE 24 experiment

Figure S4a – Average abundance coverage of the three major ion types in unmodified peptide spectra

Figure S4b – Average abundance coverage of the three major ion types in oxidized peptide spectra

Figure S4c – Average abundance coverage of the three major ion types in phosphopeptide spectra

Figure S5 – The head-to-tail plots of the predicted and experimental spectrum from the test set

Figure S6 – The head-to-tail plot of the predicted and experimental spectrum from the validation set

Figure S7 – Comparison of predicted and experimental spectra of a triply-charged peptide ion

Supporting Documents

Document S1 – Generation of in-silico spectral libraries

**Figure S1.** Model architecture schematic

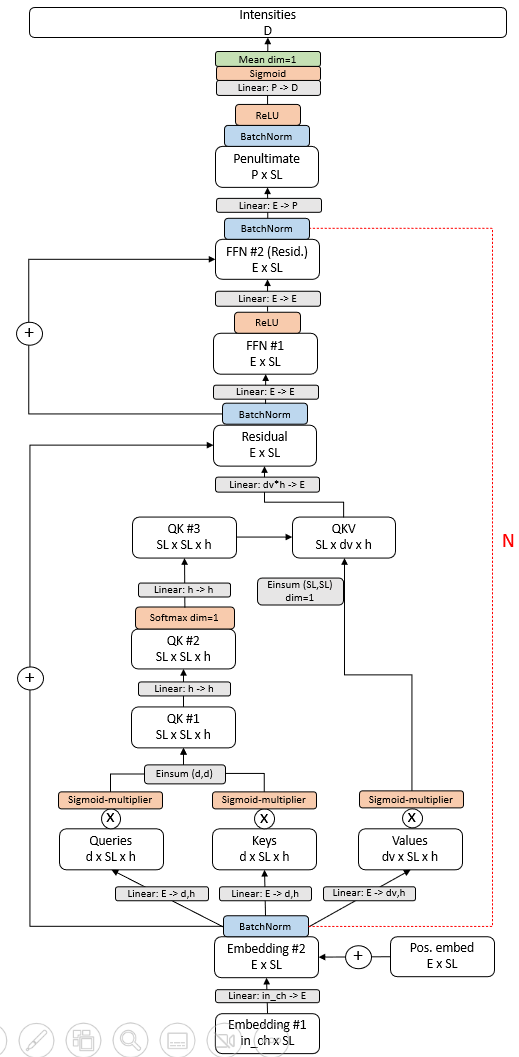

Feature layer

Linear/Einsum

Normalization

Activation

Pooling

*in_ch : Input channels (38)*

*SL : Sequence length (40)*

*E : Embed units (256)*

*d : Queries/ keys units (16)*

*dv : Values units (16)*

*h : Attention heads (64)*

*P : Penultimate units (512)*

*D : Dictionary size (7919)*

*Mean: Average pooling*

: Addition

: Multiplication

+

x

***Note:* *The detailed description of the model architecture in Figure S1.***

The model input starts at the bottom, where sequence, modifications, charge, and energy are embedded into a tensor of shape *in_ch x SL*. The *in_ch* dimension is size 38, comprised of a tiled length 21 one-hot amino acid vector, length 8 one-hot modification vector, length 8 one-hot charge vector, and a single float for collision energy in eV/100. This is done for each token/amino acid in the sequence, with max sequence length *SL* equal to 40. The precursor charge and energy values are replicated across the sequence length. The input embedding is projected to *E=*256 units, resulting in *embedding layer #2*, to which a constant learned position embedding of the same size is added. This is followed by batch normalization.

Our model is similar to that of the transformer encoder^1^, with multi-headed attention and feed forward blocks, but with minor tweaks for optimal performance on our dataset. In the attention layer, the *E x SL* input is first projected to *d=*16 with *h=*64 heads, creating the *Queries*, *Keys*, and *Values* layers. To stabilize training, each of these layers is multiplied to a single learned sigmoid activated parameter. We found that talking-heads attention^2^ worked well to make the model less sensitive to the parameter initialization. After the *Queries* and *Keys* are einsum’ed together along the *d* dimension, creating layer *QK #1*, the heads are projected to *h* new heads (note that in practice there exists a batch dimension not explicitly represented in figure S1’s notation). The resultant layer *QK #2* is softmax’ed on its second (dimension 1, zero-based) *SL* dimension, creating the attention weights. These attention weights are again projected to *h* new heads, and einsum’ed to the *Values* layer to create *dv=*16 units. Completing the attention layer is a linear transformation of the flattened *dv* and *h* dimensions to *E* units, and added as a residual to the *E x SL* input. Feed forward blocks are designed mostly unmodified from the original transformer, except that the first projection (prior to the ReLU activation) is to *E=*256 units instead of multiplying the number of units by four. Both the attention and feed forward layers are followed by batch normalization and comprise the fundamental subunit of the model. This subunit is repeated nine times.

After the transformer-like subunits, the features are projected to *P=*512 units to make the *Penultimate* layer, batch normalized, and ReLU activated. To finally make the 1D spectrum intensity predictions, the 2D *Penultimate* layer is projected to *D=*7919 (the dictionary size), a sigmoid activation to squeeze between 0-1, and mean pooled over the *SL* dimension. The m/z values must be calculated for all ion intensities in the resultant vector in order to project the prediction as a mass spectrum.

Our weight initialization scheme was ad hoc for this model, and is specific to the dimension sizes chosen in figure S1. Every linear weight is initialized to N(0, 0.03), except for *h->h* linear transformation following the softmax in the attention layer and the final linear transformation to the *Intensities*, which were initialized to N(0, 0.01).

All models were trained for 20 epochs using the Adam optimizer^3^, with each epoch being a full run through our random shuffled 1.7 million+ spectra training set. The learning rate was initially set to 3E-4 for 12 epochs and decayed by multiplying by 0.8 for each epoch thereafter. For each step the batch size is 100 spectra. The cosine similarity was scored on our validation set after each epoch, and the ultimate model used for further predictions are the weights saved from the lowest validation score.

To train all models and generate predictions/in silico libraries we used a single Dell laptop equipped with an 11^th^ generation Intel core i7-11850h @ 2.5 GHz CPU processor, and an Nvidia RTX A3000 GPU processor. The model was developed on an Anaconda 2.3.2 development platform and coded using the Pytorch 2.0.0 deep learning library.

**Figure S2.** Comparison of the CSS difference between ModelSel and ModelCons using the same peptides of the validation set. Note: ModelSel is trained by the selected library and ModelCons is trained by the consensus library. Our analysis showed that aside from the 1.7% of spectra that produced the same CSS for both models, ModelSel outperformed ModelCons in achieving higher CSS scores in 60.6% of the identifications. Conversely, ModelCons yielded superior CSS scores in 37.7% of the spectra. Given that consensus spectra are derived from peaks of many individual spectra at different collision energies, these spectra are unsuitable for training an energy-sensitive model effectively. Hence, the selected spectra were deemed most suitable for training purposes.

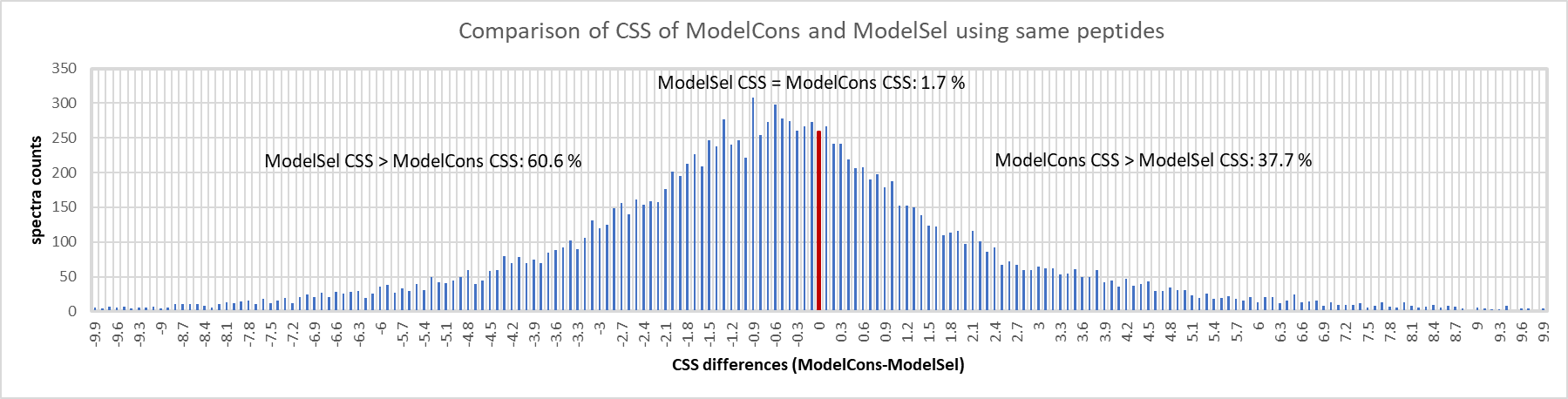

**Figure S3** **A**. Charge state errors (pointed by a red arrow) from NCE conversion to eV observed for the charge 4+ ions detected in the NISTmAb antibody NCE 24 experiment. **B**. In analyzing >8000 spectra, 18% of 2+ ions obtained deviated eVs due to incorrect ion charge state in an NCE 26 experiment.

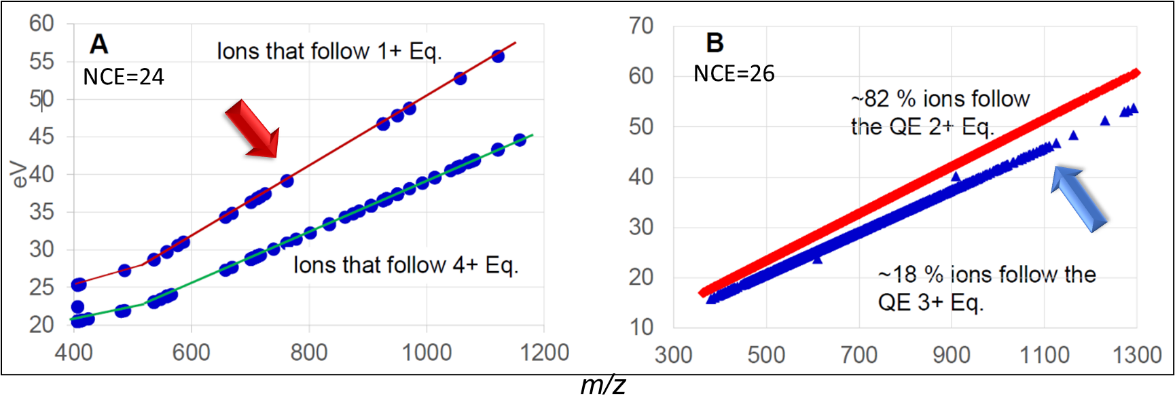

**Figure S4a**. Abundance coverage of the three major ion types in 824529 HCD spectra of unmodified peptide ions. Note: Other ions refer to neutral losses, internal ions, immonium ions, and side chain fragment ions. Unknown ions refer to potentially unannotated, contaminant, and noise peaks.

Spectra counts

| Ion abundance % per spectrum | B/y ions | Other ions | Unknown ions |
| --- | --- | --- | --- |
| 0 | 0 | 426 | 155 |
|  | 38 | 1518 | 1157 |
|  | 555 | 3340 | 4502 |
|  | 1843 | 6627 | 10128 |
|  | 3644 | 11552 | 16322 |
| 0.1 | 5432 | 18775 | 22900 |
|  | 7313 | 27941 | 29816 |
|  | 9498 | 36974 | 36137 |
|  | 11897 | 45716 | 41642 |
|  | 14484 | 53110 | 45629 |
| 0.2 | 16913 | 57818 | 48638 |
|  | 19580 | 60312 | 51182 |
|  | 22759 | 60203 | 51306 |
|  | 25620 | 58087 | 51488 |
|  | 28513 | 54554 | 50402 |
| 0.3 | 31232 | 49624 | 48832 |
|  | 33744 | 44944 | 46810 |
|  | 36311 | 39332 | 43533 |
|  | 38622 | 34355 | 40294 |
|  | 40061 | 29225 | 35960 |
| 0.4 | 40982 | 25026 | 31775 |
|  | 42357 | 21008 | 27681 |
|  | 41878 | 17023 | 23011 |
|  | 41475 | 14053 | 18440 |
|  | 39859 | 11508 | 14386 |
| 0.5 | 38397 | 9161 | 11047 |
|  | 36126 | 7226 | 7801 |
|  | 33268 | 5800 | 5274 |
|  | 30110 | 4607 | 3487 |
|  | 26656 | 3676 | 2129 |
| 0.6 | 23264 | 2727 | 1277 |
|  | 19406 | 2130 | 671 |
|  | 16002 | 1583 | 362 |
|  | 12785 | 1173 | 179 |
|  | 10078 | 857 | 74 |
| 0.7 | 7561 | 607 | 40 |
|  | 5562 | 468 | 7 |
|  | 3935 | 367 | 7 |
|  | 2653 | 264 | 2 |
|  | 1737 | 182 | 1 |
| 0.8 | 1075 | 168 | 1 |
|  | 657 | 125 | 0 |
|  | 360 | 85 | 0 |
|  | 174 | 36 | 0 |
|  | 65 | 18 | 0 |
| 0.9 | 32 | 12 | 0 |
|  | 14 | 11 | 0 |
|  | 1 | 8 | 0 |
|  | 1 | 3 | 0 |

**Figure S4b**. Abundance coverage of the three major ion types in 124723 HCD spectra of oxidized peptide ions. Note: Other ions refer to neutral losses, internal ions, iminium ions, and side chain fragment ions. Unknown ions refer to potentially unannotated, contaminant, and noise peaks.

Spectra counts

| Ion abundance % per spectrum | b/y | others | unknown |
| --- | --- | --- | --- |
| 0 | 0 | 101 | 12 |
|  | 2 | 291 | 107 |
|  | 72 | 545 | 484 |
|  | 265 | 905 | 1186 |
|  | 558 | 1406 | 2024 |
| 0.1 | 812 | 2009 | 2820 |
|  | 1264 | 2776 | 3683 |
|  | 1753 | 3626 | 4580 |
|  | 2413 | 4717 | 5431 |
|  | 3208 | 5600 | 6257 |
| 0.2 | 3932 | 6368 | 6795 |
|  | 4679 | 7065 | 7413 |
|  | 5355 | 7552 | 7680 |
|  | 6030 | 7755 | 8080 |
|  | 6352 | 8093 | 8124 |
| 0.3 | 6620 | 7661 | 7816 |
|  | 6626 | 7695 | 7598 |
|  | 6766 | 7232 | 7166 |
|  | 6632 | 6389 | 6708 |
|  | 6636 | 6179 | 6075 |
| 0.4 | 6484 | 5484 | 5409 |
|  | 6245 | 4753 | 4628 |
|  | 5897 | 4101 | 3810 |
|  | 5516 | 3536 | 3066 |
|  | 4923 | 2975 | 2407 |
| 0.5 | 4518 | 2321 | 1880 |
|  | 3999 | 1829 | 1341 |
|  | 3415 | 1489 | 837 |
|  | 2871 | 1049 | 533 |
|  | 2449 | 876 | 341 |
| 0.6 | 2086 | 594 | 198 |
|  | 1654 | 470 | 120 |
|  | 1263 | 329 | 64 |
|  | 957 | 253 | 21 |
|  | 763 | 198 | 17 |
| 0.7 | 587 | 151 | 5 |
|  | 390 | 93 | 2 |
|  | 269 | 57 | 0 |
|  | 172 | 52 | 0 |
|  | 113 | 30 | 0 |
| 0.8 | 70 | 25 | 0 |
|  | 48 | 17 | 0 |
|  | 26 | 11 | 0 |
|  | 18 | 8 | 0 |
|  | 7 | 6 | 0 |
| 0.9 | 5 | 0 | 0 |
|  | 2 | 0 | 0 |
|  | 1 | 0 | 0 |

**Figure S4c**. Abundance coverage of the three major ion types in 66918 HCD spectra of phosphopeptides ions. Note: Other ions refer to neutral losses, internal ions, iminium ions, and side chain fragment ions. Unknown ions refer to potentially unannotated, contaminant, and noise peaks.

Spectra counts

| Ion abundance % per spectrum | b/y | others | Unknown |
| --- | --- | --- | --- |
| 0 | 0 | 14 | 0 |
|  | 9 | 52 | 13 |
|  | 39 | 91 | 67 |
|  | 172 | 144 | 191 |
|  | 347 | 245 | 473 |
| 0.1 | 630 | 376 | 906 |
|  | 980 | 513 | 1509 |
|  | 1460 | 809 | 2263 |
|  | 1995 | 1117 | 3092 |
|  | 2457 | 1485 | 3706 |
| 0.2 | 2997 | 1937 | 4491 |
|  | 3486 | 2394 | 4978 |
|  | 3814 | 3052 | 5211 |
|  | 4122 | 3534 | 5250 |
|  | 4256 | 4097 | 5329 |
| 0.3 | 4410 | 4247 | 5121 |
|  | 4226 | 4511 | 4756 |
|  | 4102 | 4558 | 4209 |
|  | 3929 | 4314 | 3768 |
|  | 3782 | 4004 | 3108 |
| 0.4 | 3263 | 3637 | 2452 |
|  | 2925 | 3303 | 1869 |
|  | 2538 | 3050 | 1459 |
|  | 2244 | 2726 | 1073 |
|  | 1902 | 2315 | 666 |
| 0.5 | 1493 | 1946 | 393 |
|  | 1265 | 1722 | 266 |
|  | 954 | 1436 | 162 |
|  | 761 | 1224 | 64 |
|  | 592 | 1013 | 38 |
| 0.6 | 458 | 756 | 20 |
|  | 355 | 655 | 12 |
|  | 271 | 503 | 2 |
|  | 214 | 385 | 1 |
|  | 143 | 291 | 0 |
| 0.7 | 94 | 191 | 0 |
|  | 78 | 106 | 0 |
|  | 64 | 81 | 0 |
|  | 32 | 43 | 0 |
|  | 28 | 19 | 0 |
| 0.8 | 12 | 9 | 0 |
|  | 8 | 7 | 0 |
|  | 7 | 0 | 0 |
|  | 1 | 0 | 0 |
|  | 3 | 0 | 0 |
| 0.9 | 0 | 0 | 0 |
|  | 0 | 0 | 0 |
|  | 0 | 0 | 0 |
|  | 0 | 0 | 0 |
|  | 0 | 0 | 0 |

**Figure S5**. A comparative analysis of the predicted and experimental spectra from the test set using head-to-tail plots. A. The head-to-tail plot displays the predicted spectrum (inclusive of parent peaks) for a doubly-charged peptide ion, TVPGPLFTDFVRPLNINPDR, in contrast to its corresponding spectrum in the test set. B. A comparison of the same two spectra, excluding parent peaks, showcases only the fragment ions. This example demonstrates a low-scoring prediction resulting from particularly challenging spectra, containing unfragmented parent peaks under inefficient fragmentation energies.

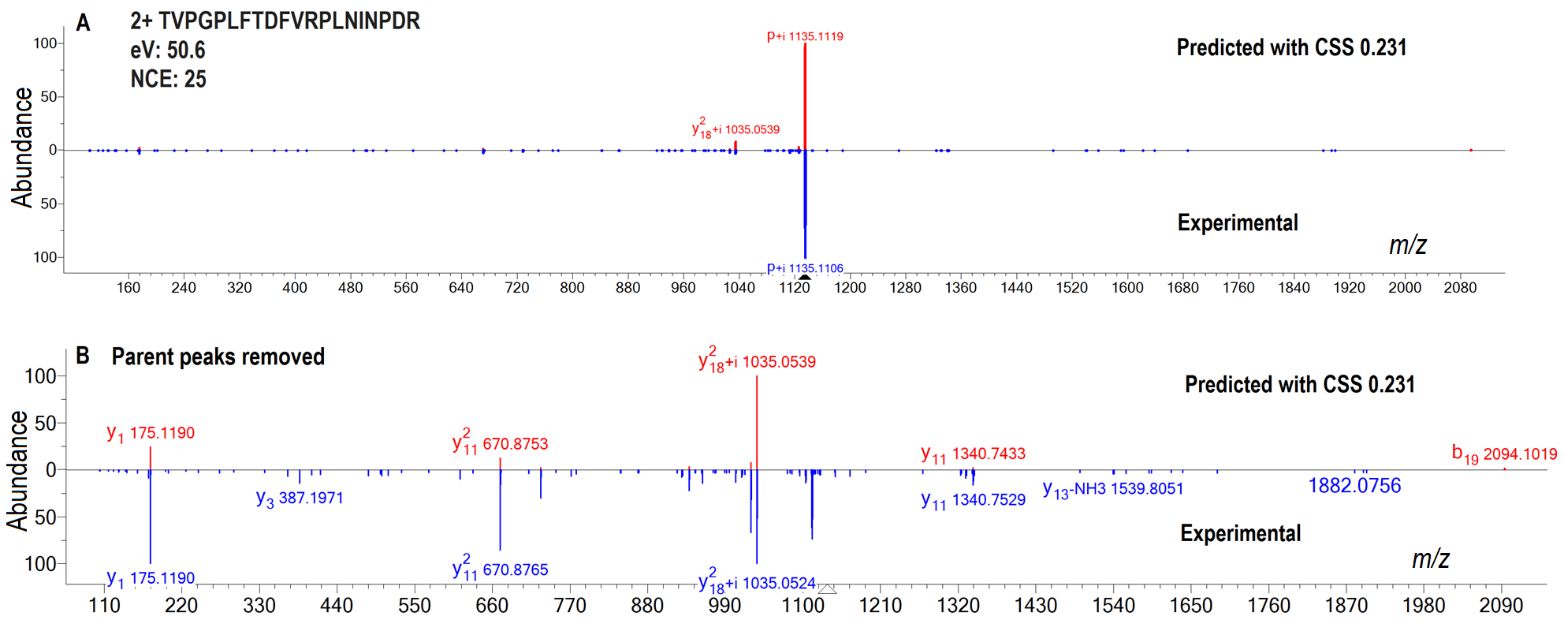

**Figure S6**. A head-to-tail plot comparing the predicted and experimental spectra from the validation set, focusing solely on fragment ions by excluding parent peaks. This example highlights a low-scoring prediction due to significantly different intensities, arising from intricate spectra of a large peptide composed of 39 amino acids.

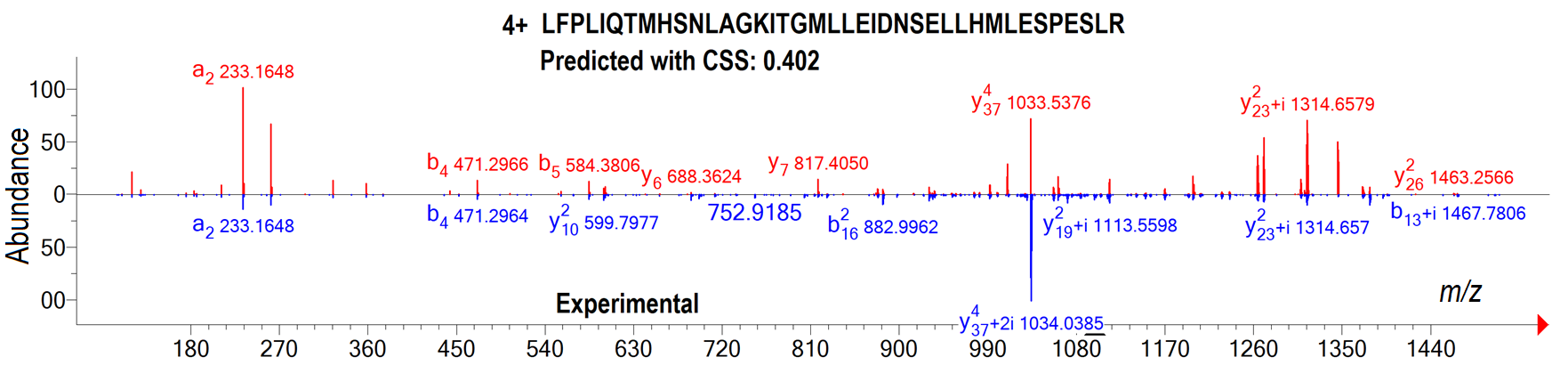

**Figure S7**. Comparison of predicted and experimental spectra of a triply-charged peptide ion with the sequence LGVNNISGIEEVNMFTNQGTVIHFNNPK are displayed in the head-to-tail plots. Plot A shows very different spectra (Prediction in the upper panel and query in the bottom panel) of this identified peptide ion. Plot B shows a good match between the same predicted spectrum (upper) and the NIST library reference spectrum (lower panel).

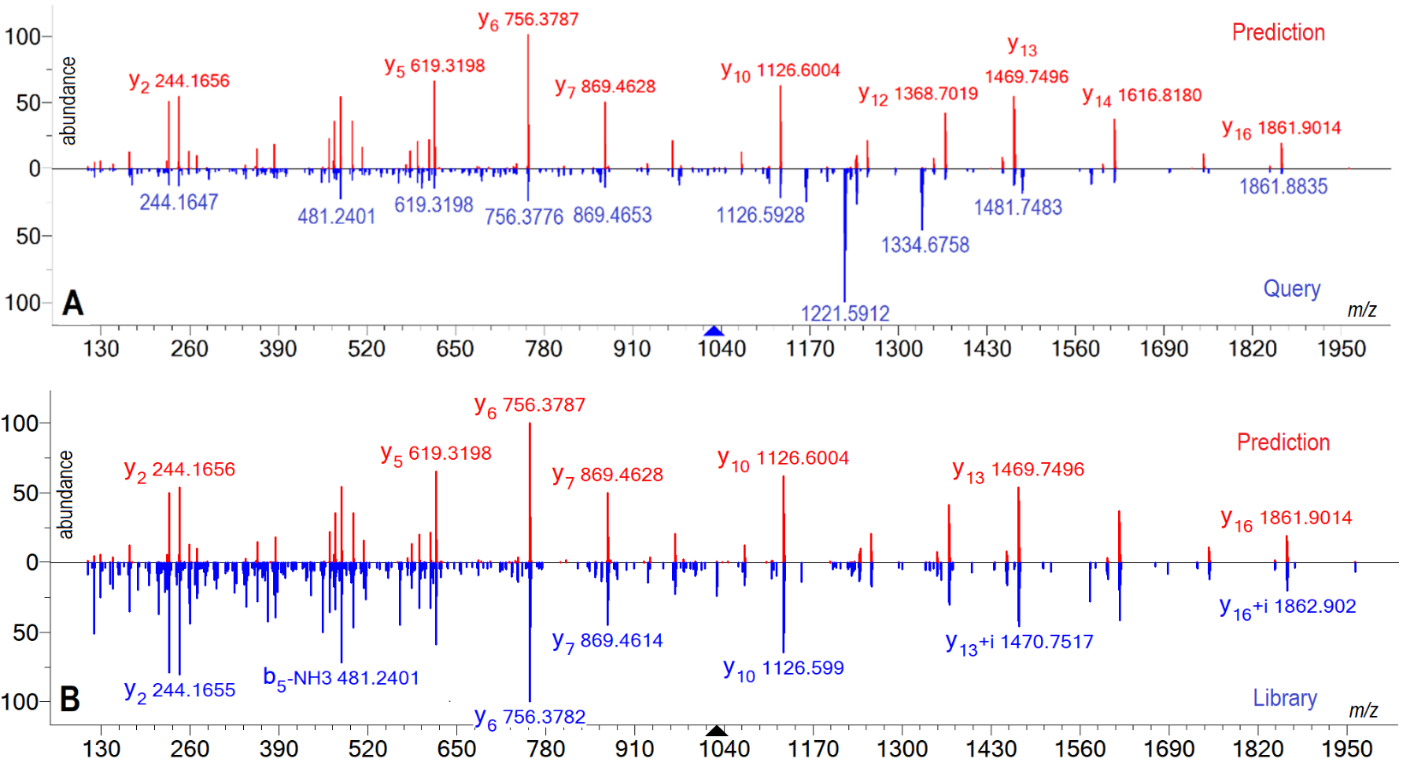

***Note: Outcomes of the three search methods display noticeable differences***

While conducting a manual assessment of peptide assignment discrepancies, specifically concentrating on high-scoring identifications exclusive to in-silico libraries but with low scores assigned or undetected by MS-GF+, we observed that when b series become predominant ions and y series are significantly diminished in these mass spectra, MS-GF+ appears to either not recognize this pattern or allocate low scores for such PSMs. Though tryptic peptides generally exhibit pronounced basicity at the C-terminus, leading to a more noticeable y-ion series, it is not uncommon for peptides with N-terminal basic residues to generate intense b-ion ladders. Through manual analysis, we determined that our model can accurately predict this kind of spectra with higher match scores.

In a similar vein, we assessed cases where MS-GF+ produced higher scores, but in-silico libraries yielded scores beneath the cutoff for peptide identifications, resulting in 3560 such identifications. Among them, we identified approximately 27% inaccurate fragment intensity predictions for certain peptides, particularly large peptides with numerous internal basic sites and proline residues as we discussed earlier in the model evaluation. Furthermore, we uncovered 73% of instances where the impact of fragmentation pattern variations can be observed on library search results. This can be illustrated by an instance of a triply-charged peptide ion with the sequence LGVNNISGIEEVNMFTNQGTVIHFNNPK and its corresponding spectra depicted in Figure S-7's head-to-tail plots. Plot A highlights the striking differences between the predicted spectra (upper panel) and query spectra (lower panel) fragmentation patterns for this peptide ion, resulting in a match score of 390 for the prediction. Conversely, Plot B exhibits an almost perfect match between the same predicted spectrum (upper) and the NIST library reference spectrum when excluding low-mass region noise (lower panel). Manual examination confirmed MS-GF+'s correct assignment of the query spectrum to this peptide ion, while also indicating that the low-scoring in-silico spectra were precisely predicted for the same peptides, as corroborated by the experimental library reference spectrum displayed in Figure B. The low scores were assigned since the query spectra deviated from previously measured spectra in the NIST spectral libraries. It is important to mention that while consistent peptide fragmentation patterns are more probable under similar collision energies and instruments, some MS/MS spectra, such as this subset, seem to exhibit significantly altered fragmentation patterns, which can influence the spectral similarity score. This finding implies that spectral matching results relying on fragment ion intensity patterns might be somewhat affected by experimental spectral reproducibility; on the other hand, database search engines that disregard fragment patterns seem to be less impacted by MS/MS reproducibility issues.

***Document S1.*** *Generation of in-silico spectral libraries*

The creation of in-silico spectral libraries for proteomics applications entailed a series of steps.

1. An in-house C++ program was used to perform in-silico tryptic digestion on the human proteome fasta file (UniProt_human_reviewed_20191015). This produced potential tryptic peptide ions with estimated charges based on the basic sites and computed their theoretical *m/z* values using specific parameters. These parameters encompassed two missed cleavages, fixed cysteine carboxymethylation, protein N-terminal acetylation, and variable modifications like methionine oxidation as well as serine, threonine, and tyrosine phosphorylation. Variable modifications were allowed to occur at any single site or all sites in the peptide.
2. The 14043412 unique peptides generated in Step 1 were filtered according to model suitability and experimental criteria (i.e., charge states ranging from 2+ to 5+, peptide length between 8 and 40, and precursor *m/z* >350), which refined the list to 5853496 peptides. An additional 394124 observed peptides from the NIST spectral library, not included in the theoretical list, were combined with this list, yielding a total of 6247620 unmodified and modified peptide ions.
3. The subsequent step involved in-silico collision energy optimization for all peptide ions, i.e., preparing their optimal energy conditions for targeted experiment-based predictions. For each ion, a chosen NCE range derived from the target experimental energy was converted into applied energies (eV). At this point, a list of spectral labels consisting of sequence, charge, modification, and collision energy eV was complete and ready for spectrum prediction.
4. a. The resulting list of spectral labels was fed through our model to predict fragment ion spectra and produce spectral libraries in MSP format. The comprehensive proteome-wide in-silico spectral library encompassed (number of NCEs) x 6247620 spectra, resulting in > 80 million spectra. The energy setting used for prediction included 13 NCEs ranging from 16, 20, 24, 28, 31, 32, 33, 35, 36, 40, 44, 48, to 52.
   b. Alternatively, a list of sequences, charges, modifications, and collision energies could be directly extracted from existing mass spectral libraries or LC-MS/MS datasets, obviating steps 1-3. Following this, model predictions reproduced these comprehensive libraries and datasets.
5. To utilize these libraries with MS PepSearch, the library’s MSP format was converted into the NIST standard library format using the Lib2NIST library conversion tool, accessible at https://chemdata.nist.gov/mass-spc/ms-search/Library_conversion_tool.html.
6. In the last step, following the prediction of target libraries, the corresponding decoy libraries were predicted on the entire target library by reversing peptide sequences, excluding the C-terminal residue, and relocating any modification along with its modified residue to the new position.

References:

1. Vaswani, A.; Shazeer, N.; Parmar, N.; Uszkoreit, J.; Jones, L.; Gomez, A.N.; Kaiser, L.; Polosukhin, I. Attention is all you need. *Adv Neural Inf Process Syst. 2017, 5998--6008.*
2. Shazeer, N.; Lan, Z.; Cheng, Y.; Ding, N.; Hou, L. Talking-Heads Attention. CoRRarXiv:2003.02436. 2020.
3. Kingma, D. P.; Ba, J. Adam; A method for stochastic optimization. Proceedings of the 3 rd International Conference on Learning Representations (ICLR). 2015
